# Arabidopsis AtPME2 has a pH-dependent processivity and control cell wall mechanical properties

**DOI:** 10.1101/2021.03.03.433777

**Authors:** Ludivine Hocq, Olivier Habrylo, Aline Voxeur, Corinne Pau-Roblot, Josip Safran, Fabien Sénéchal, Françoise Fournet, Solène Bassard, Virginie Battu, Hervé Demailly, José C. Tovar, Serge Pilard, Paulo Marcelo, Brett J. Savary, Davide Mercadante, Maria Fransiska Njo, Tom Beeckman, Arezki Boudaoud, Jérôme Pelloux, Valérie Lefebvre

## Abstract

Pectin methylesterases (PMEs) modify homogalacturonan’s (HG) chemistry and play a key role in regulating primary cell wall mechanical properties. How PME activity can fine-tune pectin structure in the growing plant has remained elusive. Here we report on the Arabidopsis AtPME2, which we found to be highly expressed during lateral root emergence and dark-grown hypocotyl elongation. We produced the mature active enzyme using heterologous expression in *Pichia pastoris* and characterized it through the use of a generic plant PME antiserum suitable for detecting recombinant and native enzyme independent of species source. At neutral pH AtPME2 is preferentially active on pectins with a degree of 55-70% methylesterification and can be inhibited by PME inhibitor protein (PMEI). We show that the mode of action for AtPME2 can switch from full processivity (at pH 8), creating large blocks of unmethylated galacturonic acid, to low processivity (at pH 5) and relate these observations to the differences in electrostatic potential of the protein at acidic and alkaline pH. To assess the role of AtPME2 in development, we characterized two knock-out lines. We show that in the context of acidified apoplast, low-processive demethylesterification by AtPME2 can loosen the cell wall, with consequent increase in cell elongation and etiolated hypocotyl length. Our study brings insights into how the pH-dependent regulation by PME activity could affect pectin structure and associated cell wall mechanical properties in expansion.

**One sentence summary:** The processivity of AtPME2, a pectin methylesterase that fine-tunes cell wall pectins is modulated by pH *in vitro* and impacts the mechanical properties of the wall, affecting development *in planta*.

## Introduction

How plants control pectin’s chemistry in cell walls is a central question in plant growth and development and in plant response to abiotic and biotic stresses. Pectins are complex polysaccharides that function as key structural elements regulating the mechanical properties of plant cell walls. Pectins are enriched in galacturonic acid and comprise four main domains: homogalacturonan (HG), rhamnogalacturonan-I (RG-I), rhamnogalacturonan-II (RG-II) and xylogalacturonan (XG). One key feature of HG chemistry, a homopolymer of α-1,4-linked-D-galacturonic acid units, is the presence of methyl- and acetyl-ester substitutions along the polymer chain that modify its physical, chemical, and biochemical properties (Ridley et al., 2001). Plants synthesize HG as a highly methylesterified form (up to 80% methyl esters, occurring at the C-6 carboxyl position) and a low acetylated form (up to 5-10% acetyl ester, occurring at the O-2 or O-3 positions) in the Golgi apparatus, before being exported to the apoplastic space. The degree of methylesterification (DM) and degree of acetylation (DA), as well as the distribution of these substitutions on the backbone are fine-tuned at the cell wall by pectin methylesterases (PMEs, EC 3.1.1.11) and pectin acetylesterases (PAE, EC 3.1.1.6), respectively (Pelloux et al., 2007). Pectin methylesterase action on HG is tightly regulated biochemically by proteinaceous inhibitors called pectin methylesterase inhibitors (PMEIs), or by pH and cations (Micheli, 2001). Resulting activity can introduce extensive de-methylesterified HG blocks that can bind Ca^2+^ ions cooperatively, creating so called “egg-box”, cross-linked structures that promote cell wall rigidity (Willats et al., 2006). Limited de-methylesterified blocks may also provide substrate-binding sites for pectin-depolymerizing enzymes such as polygalacturonases (endo-PGs, EC 3.2.1.15) and pectin/pectate lyases-like (PLLs EC 4.2.2.2), which reduce HG’s degree of polymerization (DP) and promotes the pectic network’s deconstruction (Sénéchal et al., 2014b). Therefore, to relate the consequences of PME action on pectin substrates to changes in the cell wall’s elasticity, it is key to determine their degree of processivity (*i.e*., the extent PME hydrolyzes consecutive methylesters).

Plants are well described for expressing multiple PME isoforms with individual isozymes varying in tissue-specific expression patterns, biochemical properties, and action patterns. PMEs thereby likely function differentially in the cell wall during plant growth and development (Goldberg et al., 1996; Micheli, 2001; Pelloux et al., 2007). PMEs were indeed reported to play a key role in developmental processes as diverse as hypocotyl elongation (Pelletier et al., 2010), pollen tube growth (Leroux et al., 2015), root development (Hewezi et al., 2008), organogenesis at the shoot apical meristem (Peaucelle et al., 2008; Peaucelle et al., 2011), and gynoeceum development (Andres-Robin et al., 2018). Contradictory reports showed that PME activity can either induce cell wall stiffening or loosening, with distinct consequences on plant development (Peaucelle et al., 2015; Daher et al., 2018; Wang et al., 2020). This could at least partly be explained by the demethylation pattern that different PME isoforms would create in relation to their processivity, which could be regulated by the local cell wall microenvironment, including ion concentrations, apoplastic pH, enzyme’s localization, and presence of inhibitory proteins.

Because plant PMEs are encoded by large multigenic family (e.g., 66 genes in Arabidopsis; Sénéchal, Wattier, et al., 2014), there is need to determine the expression profile and degree of processivity of individual isoforms to assess their potential for generating HG micro-domains that differ in de-methylesterified block sizes. Such micro-domains were recently reported to play a key role in determining the control of mucilage release in Arabidopsis seeds through interaction with peroxidases (Francoz et al., 2019). Plant PMEs typically have neutral to alkaline pH activity optimum (Jolie et al., 2010; Dixit et al., 2013) although few acidic isoforms are reported (Lin et al., 1989; Thonar et al., 2006). It is generally recognized that plant and microbial PMEs differ in their processivity. Plant and bacterial PMEs produce large blocks of demethylesterified HG by processive action, while fungal enzymes act more randomly on their substrate, providing single or limited consecutive demethylesterifications (Fries et al., 2007; Mercadante et al., 2013; Mercadante et al., 2014; Sénéchal et al., 2015; Kent et al., 2016). The structural determinants for differences in processivity were assessed (Mercadante et al., 2014; Kent et al., 2016), with key suggestions about the role of charged residues in certain subsites of the enzyme binding groove, and the interplay of electrostatic *versus* hydrophobic contacts in favoring substrate-binding and sliding along the groove to achieve processivity (Fries et al., 2007; Mercadante et al., 2014; Kent et al., 2016). The myriad of PME isoforms expressed in plants is however suggestive of a very fine regulation of the processive activity. The binding to certain methylation pattern and the enzymatic release after a certain number of de-methylesterification cycles are likely to be fine-tuned to modulate the physico-chemical properties of plant cell wall pectin in accordance to the micro-environment. PME processivity of an apple PME was shown to be pH-dependent, with a possible shift from a blockwise to non-blockwise mode of action (Denès et al., 2000), and processive fungal PMEs were also reported (Markoviě and Kohn, 1984; Safran et al., 2021). Considering such complexity of the PMEs’ landscape, it is therefore paramount importance to adopt a more comprehensive approach in which biochemical data are combined with structural and biophysical information of PMEs activity. Nevertheless, studying the crystal structure or mode of action of rare plant PMEs (i.e., low abundant proteins due to limited temporal and tissue-specific expression) has been impaired by the ability to produce purified native enzyme in quantities sufficient for refined structural studies. Routine heterologous expression of effectively folded plant PMEs has been challenging, thus their precise mode of action remains unresolved (Cheong et al., 2019).

We describe here the biochemical and functional characterization of Arabidopsis *AtPME2* (At1g53830), a group 2 PME (harboring a N-terminal extension, i.e. PRO-domain, showing sequence similarities with the PMEI domain (Pfam04043)) which is strongly expressed in dark-grown hypocotyls and roots, and whose protein localizes at the cell wall. We successfully expressed active AtPME2 in the yeast *Pichia pastoris*, and using generic PME antibodies generated from a designed peptide immunogen, we show that the PRO-part is important for processing the enzyme into its mature active form. We further determined that AtPME2 is more active on moderately to highly methylesterified pectic substrates, with a high processivity at neutral pH, while it shows a low degree of processivity in acidic conditions. And finally, using loss-of-function mutant plants for *AtPME2*, we show that the enzyme may play a key role in controlling dark-grown hypocotyl development through modulating the cell wall’s structural chemistry and mechanics. This study brings insights on how the differential expression of an individual Arabidopsis PME isoform having distinctive processivity for homogalacturonan may contribute to structural changes in the cell wall that affect plant development.

## Results

### *AtPME2* gene is expressed in dark-grown hypocotyls and roots

*AtPME2 (At1g53830)* gene expression was followed using RT-qPCR transcript profiling in various organs (roots, dark-grown hypocotyls, leaves, stem, siliques, floral buds and seeds) and was found to be highly expressed in dark-grown hypocotyls and roots as compared to leaves, stem, floral buds and seeds. In contrast, no expression was detected in siliques (**Figure 1A**). During the time course of dark-grown hypocotyl development, an increase in *AtPME2* transcripts was measured up to 72 h post-induction (**Figure 1B**). This timing corresponds to the acceleration phase of growth according to previously published work (Pelletier et al., 2010). In contrast, *AtPME2* was stably expressed in roots during seedling development in the light (**data not shown**).

**Figure 1:**
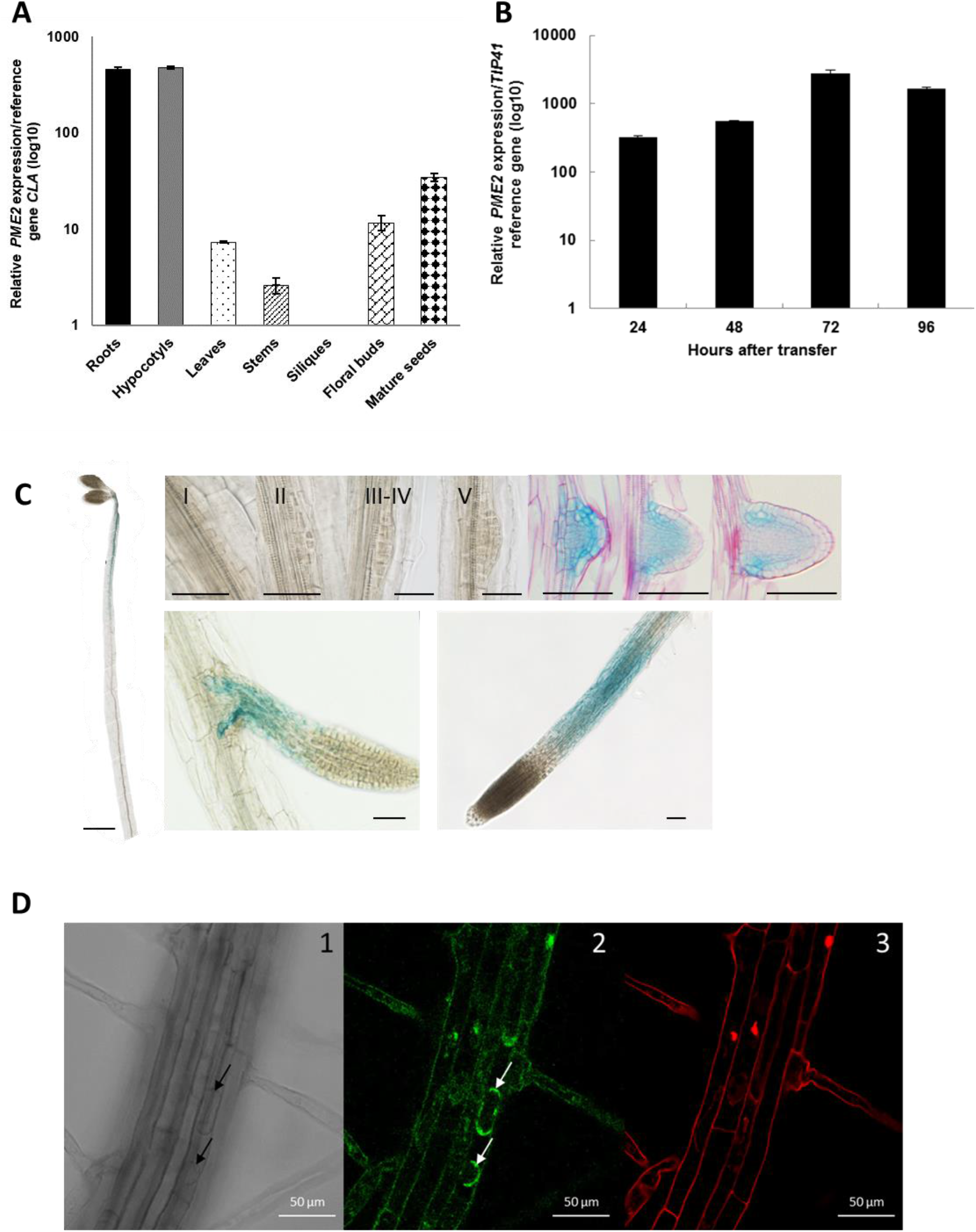
*AtPME2* gene is highly expressed in roots and dark-grown hypocotyls and targeted at the cell wall. At*PME2* gene expression was quantified **(A)** on different organs and quantified using *CLATHRIN* (*At5g46630*) as a reference gene, **(B)** on various stages of dark-grown hypocotyl elongation (up to 96h post-induction) and quantified using *TIP41* (*At4G34270*) as a reference gene and (**C**) Localization of AtPME2 promotor activity during in 4 day-old dark-grown hypocotyl (left, scale bar: 1 mm) and lateral root initiation (right, scale bars: 100 μm). (**D**) Subcellular localization of the AtPME2 protein. Arabidopsis plants were transformed with Rhizobium radiobacter containing a 35S::AtPME2-GFP construct and GFP fluorescence was imaged in 6-day old roots under confocal microscope. 1. Brightfield imaging of plasmolyzed root cells, 2. AtPME2-GFP fused protein signal, 3. Propidium iodide staining of the cell walls. Scale bar: 50 μm; Arrows indicate retraction of the tonoplast due to plasmolysis.

*AtPME2* promoter activity was further localized using a *GUS* reporter gene. Following plant transformation, GUS staining was assessed in light-grown and 4 day-old dark-grown seedlings. In etiolated hypocotyls, the promoter activity was mainly localized in the upper part of the organ (**Figure 1C, left panel**). During lateral root formation, no GUS staining was detected in the early stages (stages I to V) of primordia differentiation, while a strong signal was observed at later stages (from VI onwards) (**Figure 1C, upper panel**). In elongating roots (either primary, lateral or adventitious), *AtPME2* promoter activity was mainly present in the elongation zone (**Figure 1C, lower panel**).

### AtPME2 protein is present as a processed isoform in the cell wall

Using proteomic profiling, we identified pectin remodeling enzymes PME, PAE, PG, PLL and regulatory proteins PMEI and SBT (subtilase) in cell wall-enriched protein fractions isolated from either 4-day-old hypocotyls (**Supplemental Table IA**) and 7-day-old roots (**Supplemental Table IB**), of Col-0 and WS ecotypes. This survey confirmed that AtPME2 was indeed present in cell wall-enriched protein fractions of both organs, thus supporting transcriptional data.

To further verify the secretion of AtPME2 in the apoplasm, we designed a genetic construct tagging AtPME2 with GFP attached at the C-terminus of the mature protein sequence. Following plant transformation, confocal imaging of plasmolyzed root cells that revealed GFP fluorescence was detected at both the cell wall and the cytoplasm (**Figure 1E**). This is consistent with AtPME2 translocation to the cell wall where it acts to fine-tune pectin structure and with previous reports of the processing of nascent into mature protein during transport in Golgi vesicles (Micheli, 2001; Wolf et al., 2009).

### AtPME2 can be effectively produced and processed as an active isoform in *Pichia pastoris*

To produce pure active enzyme for biochemical studies we took into consideration the hypothesis that the PMEI domain functions as a chaperone during PME transport and processing (Micheli, 2001). We therefore inserted the full length AtPME2 coding sequence into the pPICZ αB yeast expression vector, minus the plant secretory signal peptide and STOP codon (referred as “FL” construct, **Supplemental Figure 1**) and transformed *Pichia pastoris*. PME activity was detected in concentrated supernatants of induced transformants and recombinant AtPME2 purified after cation exchange chromatography as shown by SDS-PAGE (Coomassie-Blue stained gel) in **Figure 2A**. One band is present at ~30 kDa and two bands are observed at ~35 kDa, the latter corresponding to the approximate mass calculated from the sequence for the mature AtPME2 protein. The doublet is consistent with AtPME2 being cleaved at either of the processing motifs (RKLK and RRLL, see **Supplemental Figure 1**) by *Pichia pastoris* subtilisin protease. The identity of AtPME2 was confirmed by mass spectrometry of the tryptic peptides, matching 12 peptides of the mature protein (**Supplemental Figure 2**). The lower band, ~30 kDa, corresponds to the PRO-peptide, which was confirmed by matching 7 tryptic peptides (**Supplemental Figure 2**). Using Pichia expression system, we were thus able to produce the mature active AtPME2 enzyme, as well as recover the PRO-peptide.

**Figure 2:**
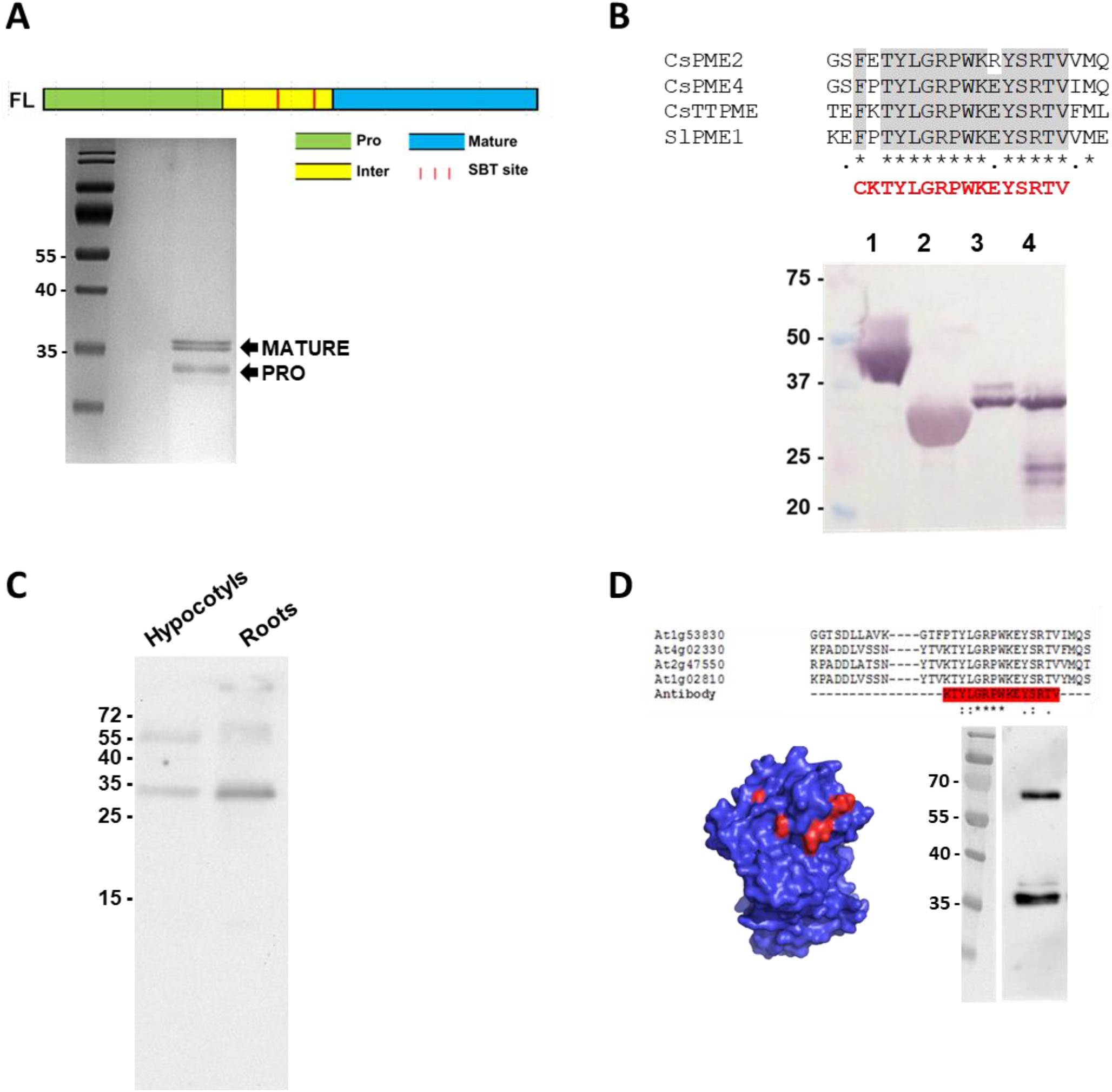
AtPME2 is effectively produced in *Pichia pastoris*. (**A**) *At*PME2 mature protein recovered from *Pichia pastoris* culture supernatant (purified by ion exchange chromatography) was separated by SDS-PAGE (Coomassie-Blue stained gel). Closely related bands at a MW ~35 kDa represent the two forms of processed enzymes (see scheme of protein structure above, including processing motifs). The lower band represent the PRO part. **(B)** Design of a peptide antibody that can detect sweet orange and tomato PME isoforms. The generic anti PME antibody was designed on a highly conserved part of the mature protein (see alignment). Western blot analysis allowed detection of purified PMEs, 1: CsTT-PME (*Citrus sinensis* thermally-tolerant isozyme; (Savary et al., 2013)), 2: CsPME2 (*C. sinensis* fruit-specific salt-independent isozyme, (Savary et al., 2010), 3: CsPME4 (*C. sinensis* salt-dependent isozyme, (Savary et al., 2010)), and SlPME1 (*Solanum lycopersicum* isozyme (Savary, 2001). **(C)** Western blot analysis of cell-wall-enriched protein extracts from 7 day-old roots and 4 day-old dark grown hypocotyls using the anti PME antibody. Both processed and non-processed forms of PME can be detected. **(D)** Western blot detection of AtPME2 purified by cation-exchange chromatography from concentrated *Pichia* culture media.

To support detection and identification of plant PMEs such as AtPME2 in expression studies, we produced an antiserum that could be broadly selective for plant PMEs (ie generic for plants, independent of species source) in western blotting. Considering the sequences alignment of 45 Arabidopsis PME catalytic domains (**Supplemental Figure 3A**), we chose a 15 amino acid peptide sequence (KTYLGRPWKEYSRTV) which is indeed highly conserved, notably in AtPME2 (AT1G53830, PTYLGRPWKEYSRTV) and AtPME3 (AT3G14310, PTYLGRPWKEYS**Q**TV) sequences, two of the proteins identified both in hypocotyls and roots in our proteomic analyses. Western blot analyses, used to assess this generic PME antibody, showed a strong antiserum binding signal for purified PME isoforms isolated from citrus (CsTT-PME, CsPME2 and CsPME4) and tomato (SlPME1) fruit (Savary, 2001; Savary et al., 2010; Savary et al., 2013) as well as purified AtPME3 (Sénéchal et al., 2015) (**Figure 2B** and **Supplemental Figure 3B**), thus supporting the generic antiserum provides a new tool for analyzing plant and Arabidopsis PMEs. To assess its sensitivity for detecting PMEs present in cell wall-enriched protein fractions, we used it to analyze Arabidopsis hypocotyl and root protein extracts. We detected antigen signals at approximately 35 kDa, which is consistent with the predicted size of fully processed (mature) PME (**Figure 2C**). Additional bands were detected above ~55 kDa, which may represent unprocessed PME precursors. Finally, we performed western-blot analysis on the recombinant purified AtPME2, and detected the two AtPME2 protein bands separated at molecular mass ~35 kDa (**Figure 2D**). A strong antigen band is also revealed with mass approaching 70 kDa, while no corresponding protein was observed in the stained protein gel (**Figure 2A**). We speculate this represents low amounts of either glycosylated unprocessed AtPME2 protein as observed in the hypocotyl and root immunoblots (**Figure 2C**), or possibly dimers formed during electrophoresis.

### Biochemical characterization of AtPME2

The pH-dependency and sensitivity to inhibition by PMEI were determined for the purified AtPME2. Using the ruthenium red gel diffusion assay with high-DM pectins (> 85%) as a substrate, we showed that the enzyme was the most active at neutral pH (7.5), although it was still active at pH 5 (**Figure 3A**). AtPME2 activity was inhibited at the three pHs tested (5, 6.3 and 7.5) by the previously reported pH-insensitive AtPMEI9 inhibitor protein (Hocq et al., 2017b). This inhibition was positively correlated with increasing quantities of AtPMEI9 (**Figure 3A**). In addition, AtPME2 could also be inhibited by pH-sensitive AtPMEI4 at pH 5 (**data not shown**).

**Figure 3:**
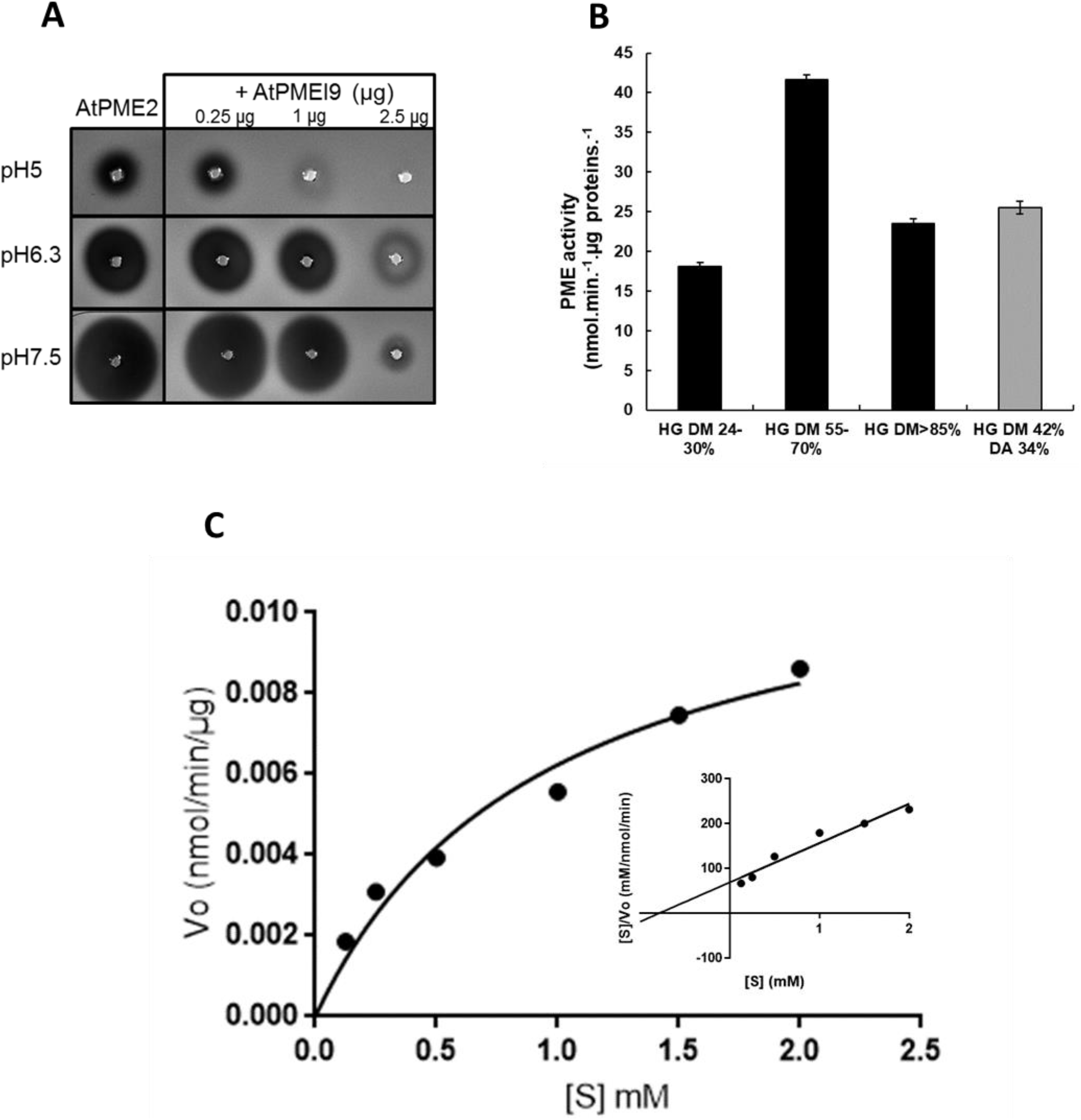
AtPME2 is active and can be inhibited by PMEI. (**A**) pH-dependence of AtPME2 activity. Activity of purified AtPME2 was assessed at three distinct pH (5, 6.3 and 7.5) with increasing quantities of the pH-independent AtPMEI9. Activity was determined with the gel diffusion assay using pectins DM 85% as a substrate and ruthenium red staining. The diameter of the halo reflects PME activity. (**B**) Substrate specificity of AtPME2. Activity of purified AtPME2 on pectic substrates with increasing degree of methylesterification (DM) was determined at an optimal pH of 7.5. Data represent the mean ± SE of three to five replicates. HG: Homogalacturonan, DA: Degree of acetylation. (**C**) Determination of Km and Vmax for AtPME2. Activity was assessed using various concentrations of pectins DM 55-10% at 37°C and pH 7.5.

We assessed AtPME2 activity on citrus pectins with varying degrees of esterification at the optimal pH 7.5. When using pectic substrates of low DA, AtPME2 activity was the strongest for DM 55 to 70% (40 nmol MeOH.min^−1^. μg proteins^−1^), and activity was reduced by ~half when using substrates of high (>85%) or low (24-30%) DM. AtPME2 was active on sugar beet pectins of DM 42% and DA 34%, suggesting acetylation of GalA residues may minimally affect the enzyme’s activity (**Figure 3B**). Using the best substrate (pectins DM 55% − 70%) and the optimal pH of 7.5, we determined the kinetic parameters of the enzyme and showed that the Km was 0.481 mM and the Vmax was 0.019 nmol MeOH.min^−1^.μg protein^−1^ (**Figure 3C**).

### AtPME2 has a low degree of processivity in acidic conditions

In order to get precise insights into AtPME2 activity, we designed an experimental set-up to characterize its degree of processivity. We first digested pectins of DM 55 to 70% with a fungal polygalacturonase (endo-PG from *Aspergillus aculeatus*) for 3 h to generate a population of HG oligogalacturonides (OGs) of various DP and DM. Following heat denaturation of PG activity, the processivity of AtPME2 was determined by characterizing the relative proportion of the resulting OGs, classified by DP and DM, after overnight incubation with 20 nmol AtPME2 at pH 8 and 80 nmol AtPME2 at pH 5 (to compensate for the lower PME activity at acidic pH). As a control, we used the commercially available PMEs extracted from Citrus peels, which also present stronger activity at neutral pH compared to acidic pH (**data not shown**). The population of OG identified after PME digestion was then compared to that obtained in non PME-treated condition (named hereafter control pH 8 and control pH 5). In control samples, we were able to detect different methylated forms for each DP, with a peak of relative abundance corresponding on average to slightly more than 50% DM (*e.g.* GalA7Me4, GalA8Me4, GalA10Me6), in accordance with the mean DM of the pectins used as a substrate. At optimal pH 8, when considering oligos of DP comprised between 3 and 10, for instance DP 10 (GalA10), the different methylated forms detected in the control samples (GalA10Me5, GalA10Me6, GalA10Me7) were totally absent in AtPME2-treated OGs. In contrast unmethylated OG trimers and tetramers were the predominant end-products that accumulated (**Figure 4A**). These OGs can result from the residual activity of Aspergillus PG in the reaction mixture: in absence of calcium *in vitro*, long blocks of unmethylated HG created by processive demethylesterification of pectins at pH8 are preferential substrates, leading to hydrolysis of the OGs pool. Similar results were obtained when using Citrus PME (**Supplemental Figure 4A)**. When PME treatment (AtPME2 and CsPME) was performed at pH 5, results were strikingly different. For a given DP, the proportion of highly methylesterified forms (> 50% DM) decreased in the PME-treated samples compared to control pH 5 (**Figure 4B, Supplemental Figure 4B**). OGs with lower DM appeared following treatment with PME (GalA10Me, GalA10Me2 and GalA10Me3, absent in control pH 5), and relative amount of higher DM decreased (GalA10Me5, GalA10Me6 and GalA10Me7) (**Figure 4B inset, Supplemental Figure 4B inset**). Results were similar for OGs of distinct DPs, including GalA6, GalA7, GalA8 and GalA9, with a shift in the abundance from highly methylesterified to low methylesterified forms of these OGs in PME-treated samples as compared to control pH5. These OGs of decreased DM are likely to have less affinity for residual Aspergillus PG as they were not further digested (**Figure 4B**). It has to be mentioned that those differences in the DM distributions are more pronounced when using AtPME2 compared to CsPME. For OGs of DP < 5, no differences were detected between samples, suggesting that PME2 and CsPME have a strong preference for substrates of DP > 5 at pH 5. Taken together, these results show that the degree of processivity of the two PMEs increases as the pH shifts from acidic towards neutral to alkaline pH.

**Figure 4:**
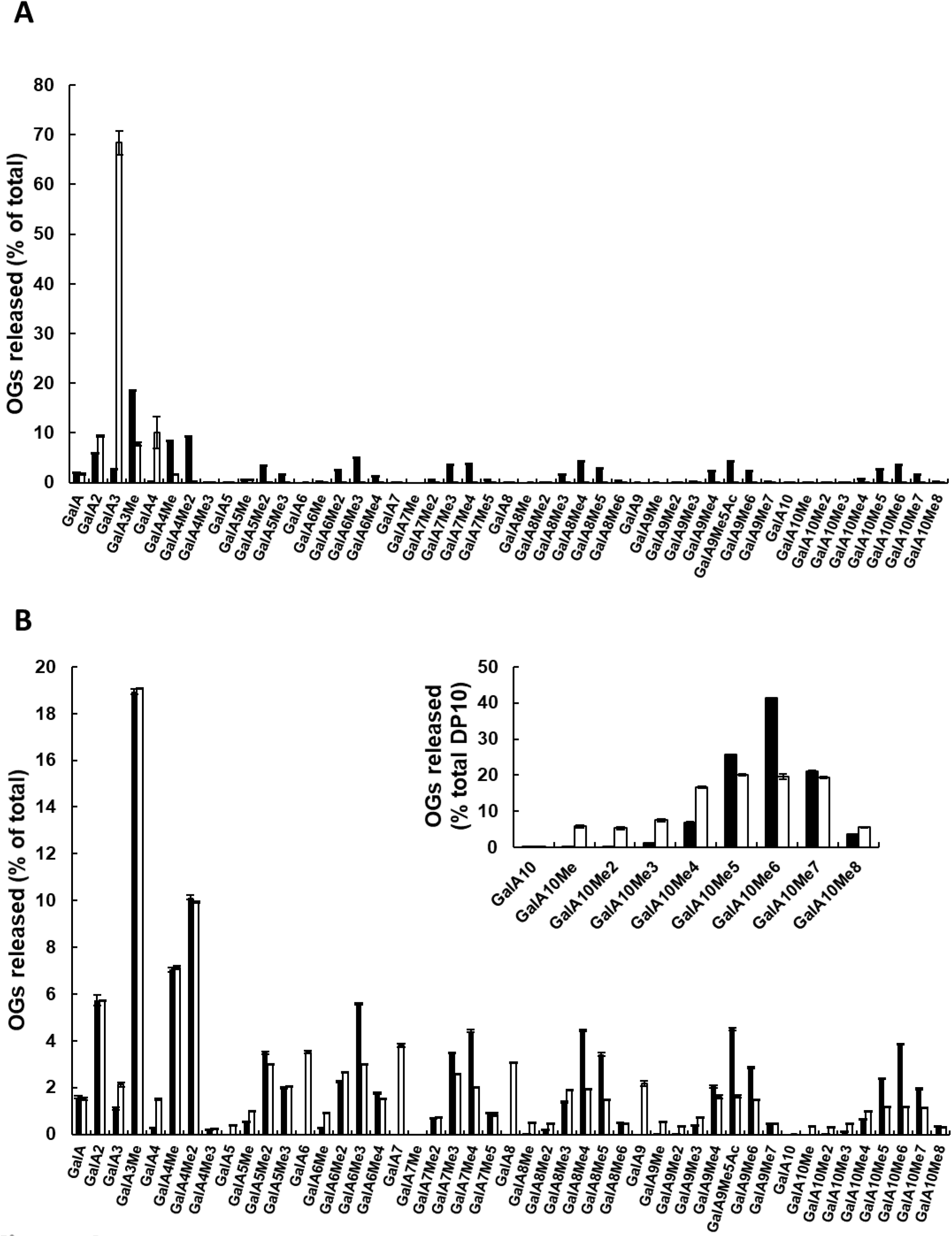
Determination of AtPME2 mode of action using an LC-MS/MS. Comparisons of the oligogalacturonides (OGs) produced following demethylesterification by AtPME2 at (**A**) pH 8 and (**B**) pH 5. A population of OGs of various degree of polymerization (DP) and degree of methylesterification (DM) was first generated by action of *Aspergillus aculeatus* polygalacturonase during 2 h at 40°C. After heat denaturation of the PGs, the OGs were incubated overnight at 40°C with buffer (black bars) or isoactivities of AtPME2 at pH 5 and pH 8 (white bars). OGs were separated using SEC and analyzed using MS/MS. Data represent the mean ± SE of three replicates.

### The electrostatic potential of PMEs correlates with their processivity

Electrostatic properties have been identified as significant in order to rationalize the basis of PMEs processivity, with charge asymmetry along the binding groove being an important feature to promote the sliding of negatively charged, demethylesterified, polysaccharides (Mercadante et al., 2014). We chose to compare the electrostatic potentials of the 2 PMEs whose modes of action were determined in this study, in addition to a fungal acidic PME from *Aspergillus niger*, AnPME, whose random mode of action at both pH has already been published (Duvetter et al., 2006; Cameron et al., 2008; Kent et al., 2016). Interestingly, AtPME2 and CsPME4 (the major isoform in the commercial PME from orange peel), which experimentally increase in processivity with increasing pH, show the largest differences in the electrostatic similarity indices, whereas AnPME, which has been indicated as a non-processive, acidic PME (Kent et al., 2016) shows little differences across pH (**Figure 5A)**. Moreover, the projection of the electrostatic potential differences between acidic and alkaline pH on the protein surfaces, normalized to highlight the differences between AtPME2 and CsPME4 or AnPME, show a concentrated positive charge patterning in the binding groove, with the largest difference observed for CsPME4 and small to no difference (electrostatic potential difference close to 0) for AnPME, as expected from comparing the action of these PMEs experimentally **(Figure 5B)**.

**Figure 5:**
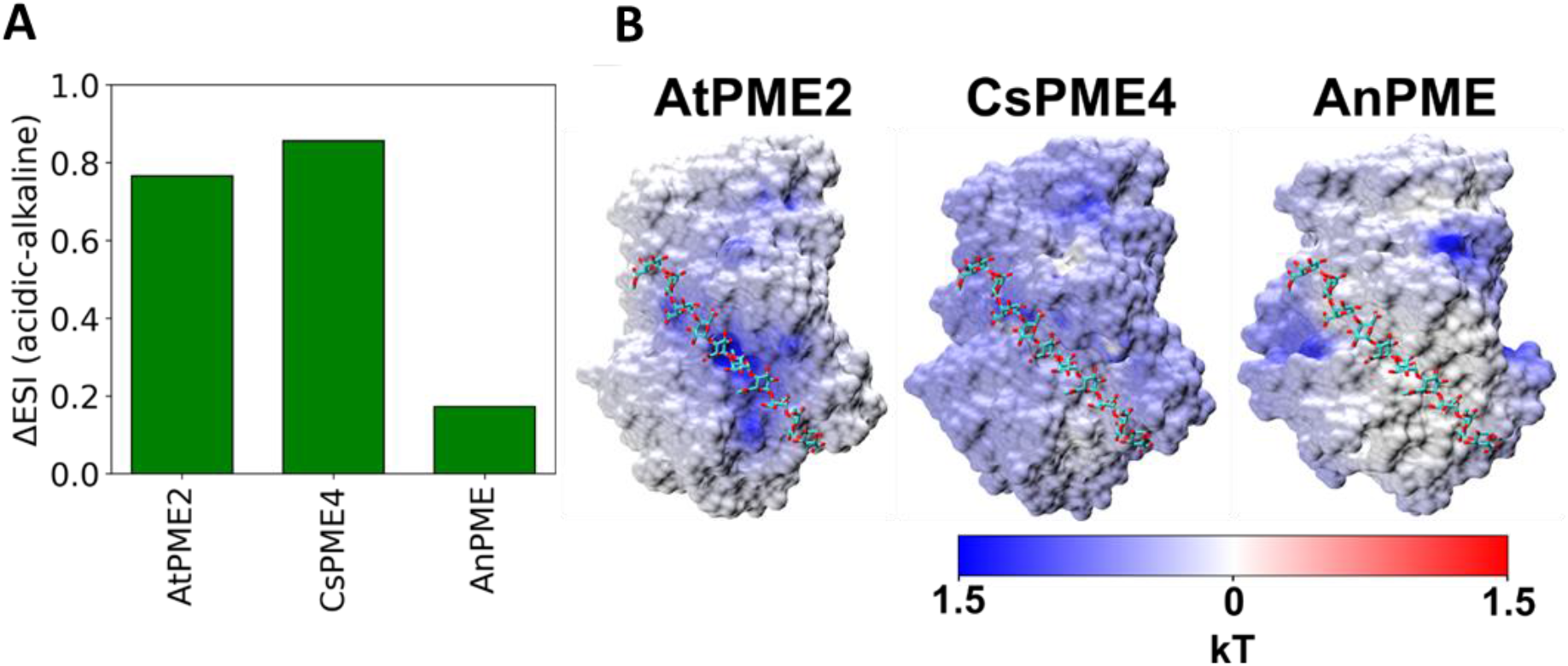
Electrostatic potential of AtPME2 is pH-dependent. **(A)** Difference between the electrostatic similarity indices of AtPME2, CsPME4 and AnPME at pH 5.0 (acidic) and pH 8.0 (basic). **(B)** Electrostatic potentials of the three PME isoforms projected on the protein surfaces. The electrostatic potentials are the resultant of the subtraction between the electrostatic potentials obtained at pH 8.0 from the one obtained at pH 5.0 for each protein. The potentials of AnPME and CsPME4 have been then divided by the electrostatic potential of AtPME2 to better show the comparison with AtPME2.

### Loss of function in AtPME2 mutants can alter pectin remodeling enzyme activities

To investigate AtPME2’s role in controlling growth and development, two homozygous T-DNA insertional knockout (KO) lines, *pme2-1* (GK-835A09, in the third exon) and *pme2-2* (FLAG_445B05, in the first exon), were identified in Arabidopsis Col-0 and WS backgrounds respectively. RT-PCR analyses revealed that both mutant lines were KO at the transcript level (**Figure 6A**). Consistent with this, no tryptic peptides from the catalytic domain of AtPME2 were detected by proteomic analyses performed on cell wall-enriched protein fractions from *pme2-1* and *pme2-2* hypocotyls (**Supplemental Table IA**). This ultimately shows that *pme2* allelic mutants are KO at the protein level. This was further supported by zymogram analysis where cell wall-enriched protein extracts from wild-type, *pme2-1* and *pme2-2* hypocotyls were resolved by isoelectric focusing (IEF) coupled with detection of PME activity. Results obtained showed no activity band at a pI of ~9 in the KO lines, which corresponds to the predicted pI of the mature part of AtPME2 (**Figure 6B**). No changes in the activity of the other PME isoforms were apparent, suggesting that the absence of the AtPME2 protein is likely to impact total PME activity. This was further confirmed by measuring pectin remodeling enzymes activities of cell wall-enriched proteins of 4 day-old dark-grown hypocotyls. As anticipated, we observed that the total PME activity decreased in *pme2* mutants compared to wild-type (**Figure 6C**). Total PG activity measured in the same type of extracts was also reduced by 10 % and 20 % in *pme2-1* and *pme2-2*, respectively (**Figure 6D**). In roots, total PME and PG activities were as well decreased in *pme2* mutants, albeit to a lesser extent compared to what was observed in dark-grown hypocotyls (**Supplemental Figure 5A and 5B**).

**Figure 6:**
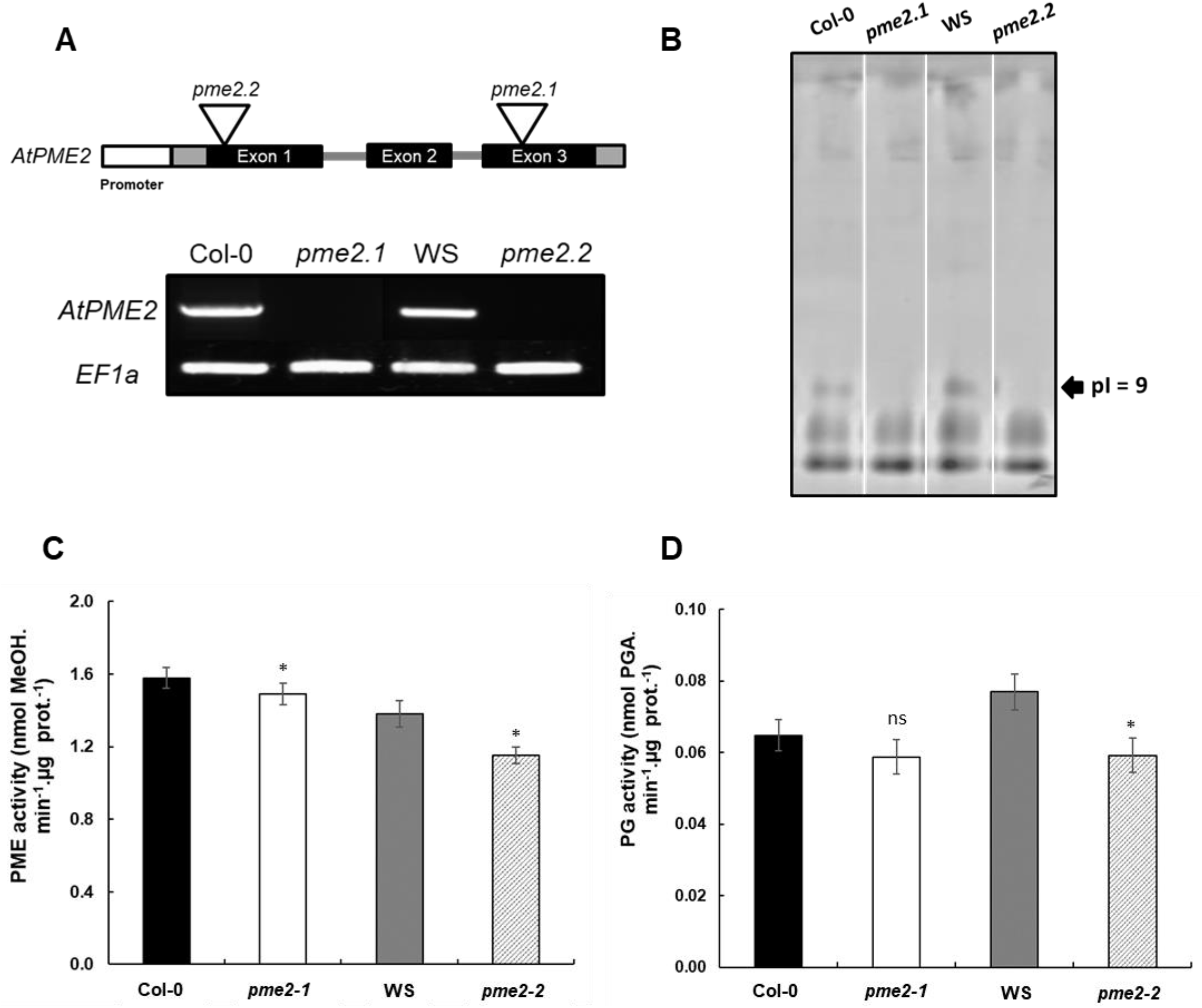
Defect in AtPME2 leads changes in pectin remodeling enzyme activities. **(A)** Schematic representation of *AtPME2* gene structure and localisation of the T-DNA insertions for *pme2-1* (GK-835A09, in the third exon) and *pme2-2* (FLAG_445B05, in the first exon). PCR analysis of *pme2-1*, *pme2-2* and WT (Col-0 and WS) hypocotyl cDNAs using specific primers flanking the T-DNA insertion sites. *EF1*◻ was used as reference gene. (**B**) Isoelectric focusing (IEF)□ of cell wall-enriched protein extracts from 4 day-old dark-grown hypocotyls of wild type Col-0/WS and *pme2-1/pme2-2* mutants. The same PME activities (15 mU) were loaded for each genotype. PME isoforms were separated and zymogram of PME activity was performed by incubating the gel with pectins (DM > 85 %), followed by ruthenium red staining. Similar observations were obtained from two independent experiments. Black arrow indicates the disappearance of an activity at a pI ~9 in *pme2* mutants. (**C**) Total PME activity of cell wall-enriched protein extracts from 4 day-old dark-grown hypocotyls of wild type Col-0/WS and *pme2-1/pme2-2* mutants. Data represent the means of PME activity in nmol of methanol.min^−1^/μg of protein^−1^± SE of three independent protein extractions and three technical replicates (n=9). (**D**) Total PG activity of cell wall-enriched protein extracts from 4 day-old dark-grown hypocotyls of wild type Col-0/WS and *pme2-1/pme2-2* mutants. Data represent the means of PG activity in nmol of PGA.min^−1^/μg of protein^−1^ ± SE of three independent protein extractions and three technical replicates (n=9). Significant differences (p<0.05*) were determined according to Wilcoxon test. Non-significant differences are indicated with ns.

### AtPME2 plays a role in controlling hypocotyl elongation through regulation of mechanical properties

To determine if changes in pectin remodeling enzyme activities may affect seedling development, we first assessed the effects of the mutations on primary root length and lateral root emergence. In our experimental conditions, elongation of primary root was slightly impaired in both mutants, although only significantly in Col-0 background, in relation to the decrease in the length of fully elongated cells (**Supplemental Figure 5C, data not shown)**. Considering AtPME2 expression pattern, lateral root density was also assessed and results showed significant lower density only for Col-0 allele (**Supplemental Figure 5D**).

We next followed etiolated hypocotyl elongation over a time-course. The rationale for etiolated hypocotyls includes: i) In the hypocotyl, cell length increased in an acropetal wave starting approximately 48 hours after sowing, ii) AtPME2 is highly expressed in the upper part of a 4 day-old growing hypocotyl, below the hook, where cells are strongly elongating, so that the absence of AtPME2 in mutants should alter their development. Thus, we hypothesized KO mutants lacking AtPME2 activity will show altered hypocotyl development. Kinematic analysis showed hypocotyl length was significantly different in both alleles, with a reduction of 10% as compared to wild-type (**Figure 7A**). The differences between wild-type and *pme2* mutants were more important from 72 h onwards, which corresponds to the rapid elongation phase. To assess if the decrease in the length is related to changes in the mechanical properties of the cell wall, we measured the stiffness (as apparent Young’s modulus) of the cell wall using atomic force microscopy (AFM, **Supplemental Figure 6**) in the *pme2-1* mutant and its corresponding wild-type Col-0. The stiffness of the epidermal cell wall at the basal part of 4-day-old hypocotyls, was similar between *pme2* and WT Col-0 (**Figure 7B, bottom panel**). In contrast, the apical part of hypocotyls (the zone where the promoter of *AtPME2* was shown to be active) showed a 10% increase in cell wall stiffness in the *pme2-1* mutant, compared to wild type (**Figure 7B, top panel**). As such, a more rigid cell wall in the *pme2-1* mutant could restrict hypocotyl elongation. The Young’s modulus measured at the top of dark-grown hypocotyl was lower (~20 MPa) than that measured in the basal part (60 MPa), also reflecting the difference of cell wall stiffness in relation to growth rate.

**Figure 7:**
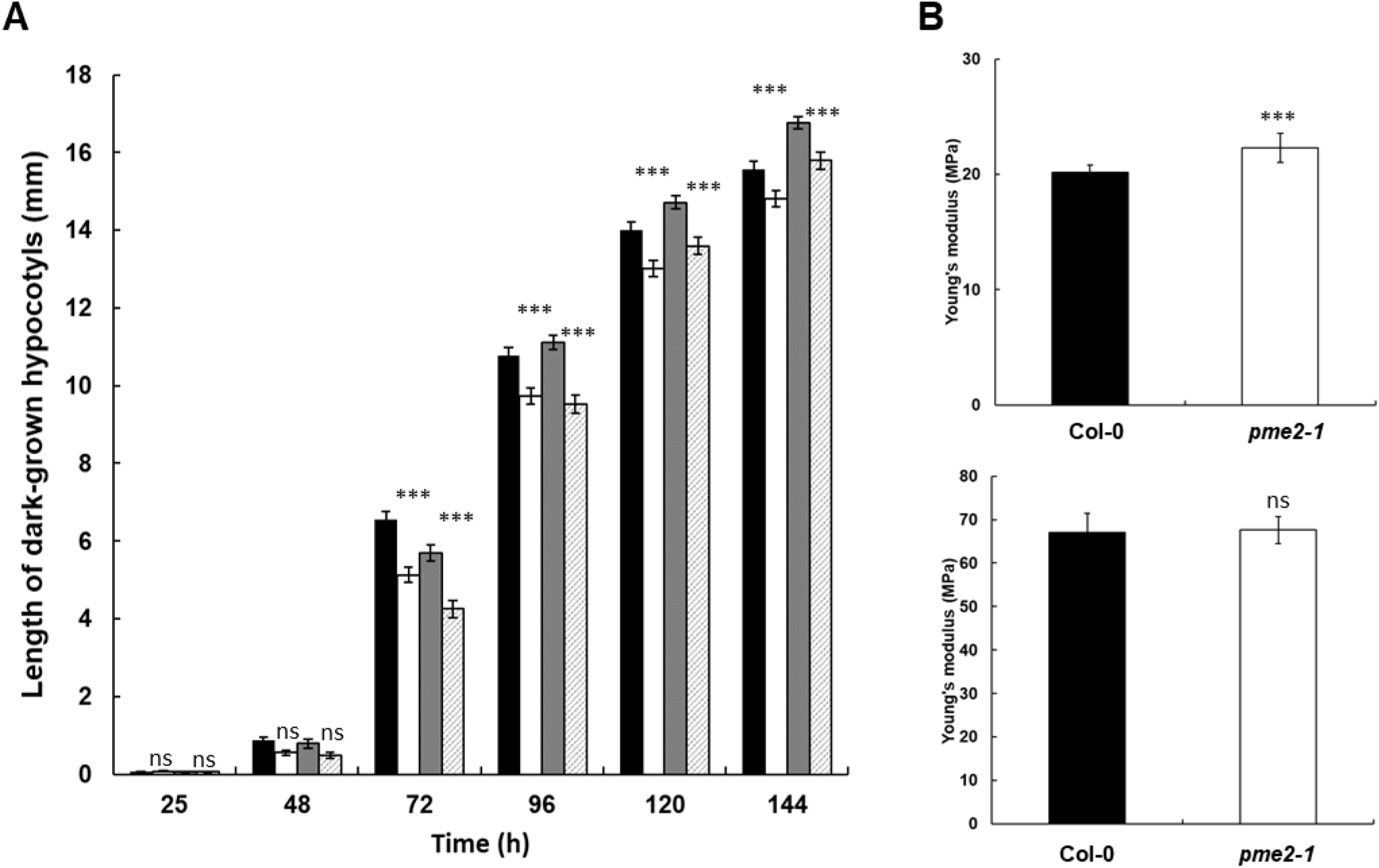
Defect in AtPME2 leads increased cell wall stiffness and reduced hypocotyl length. (**A**) Growth kinematic analysis of etiolated hypocotyls of wild type Col-0 (black bar), WS (grey bar), *pme2-1* (white bar) and *pme2-2* (hatched bar). Data represent the means of length in mm ± SE (n > 30) for each condition. (**B**) Cell wall stiffness of Col-0 (black bar) and pme2-1 (white bar) assessed by Atomic Force Microscopy on 3 day-old dark-grown hypocotyls at the bottom part (bottom panel) and below the hook (top panel). Significant differences (p<0.001***) were determined according to Wilcoxon test. Non-significant differences are indicated with ns.

## Discussion

How the mode of action by individual pectin methylesterases affect pectin’s chemistry and the mechanical properties of plant cell walls remains unresolved. This is in part due to the difficulty of obtaining purified single isoforms from plant material and to co-expression of multiple isoforms from this very large gene family. Here, we report on AtPME2, a PME expressed during lateral root emergence and in dark-grown hypocotyl elongation. Starting from its production in heterologous system and full biochemical characterization, we describe how its mode of action varies as a function of pH and assess how this might control plant development.

While *AtPME2* gene was previously found to be highly expressed during the growth transition phase in dark-grown hypocotyls (Pelletier et al., 2010), we show that, in 3 to 4 day-old seedlings, the *AtPME2* promoter is specifically active at the top of hypocotyls (**Figure 1**), a region with enhanced elongation at this stage (Refrégier et al., 2004; Peaucelle et al., 2015; Daher et al., 2018). The *AtPME2* promoter is also highly active in the root elongation zone, and during lateral root formation, as suggested by previous data sets (Brady et al., 2007; Hruz et al., 2008) and confirmed using RNA sequencing and immunocytochemistry (Wachsman et al., 2020). In lateral root primordia, the promoter’s activity initiates at stage VI before emergence (Malamy and Benfey, 1997), suggesting a role for AtPME2, together with other pectin remodeling enzymes (including PG) in the process of lateral rhizogenesis (Swarup et al., 2008; Kumpf et al., 2013; Hocq et al., 2020). Our findings are in accordance with this hypothesis (**Supplemental Figure 5**), and are further backed-up by a recent study showing that AtPME2 is of prime importance for determining lateral root mergence.

Our results support SBT-mediated processing of PME occurs in the cell before their deposition at the apoplast. Using C-terminal translational fusion, we showed AtPME2 to be present both at the cell wall and in the cytoplasm (**Figure 1A**). Cell wall-associated proteome fingerprinting analyses only identified peptides associated to mature AtPME2 and other PME isoforms from dark-grown hypocotyls and radicles in seedling (**Supplemental table IA, B**). Similar results obtained for other plant PME, in several organs, indicates thus an ubiquitous processing mechanism (San Clemente and Jamet, 2015; Sénéchal et al., 2015; Hervé et al., 2016; Nguyen-Kim et al., 2016).

Using heterologous expression to obtaining sufficient amounts of plant PME for biochemical analysis has often turned out to be challenging, despite early reports showing that AtPME31 and AtPME12 can be expressed in *E. Coli* (Dedeurwaerder et al., 2009; Cheong et al., 2019), or that an acidic PME from Jelly fig can be produced in yeast (Peng et al., 2005). While we initially failed to produce mature AtPME2 (group 2 plant PME) in *Pichia*, we subsequently showed that the PRO domain was required for proper cleavage and release of a functional PME. Our results support the hypothesis that the PRO domain supports recognition of processing motifs, RKLK and RRLL in AtPME2, by endogenous yeast subtilisins, including KEX2 and SUB2 (Bader et al., 2008; Salamin et al., 2010). *In planta*, PME and SBTs are co-expressed during development and the S1P-mediated processing of group 2 PMEs is required for the export of active enzymes in the cell wall (Wolf et al., 2009). It was further suggested that the PRO-region inhibits group 2 PMEs activity during transport through the secretory pathway (Bosch, 2005; Bosch and Hepler, 2005; Dorokhov et al., 2006). The approaches and tools that we have developed now open the way to deciphering the interaction of PME with SBT.

We found that AtPME2 has an optimal activity at weakly alkaline pH (**Figure 3A**), like Arabidopsis AtPME3 and AtPME31 (Dedeurwaerder et al., 2009; Sénéchal et al., 2015). The predicted pI~9 for AtPME2 might explain the pH-dependency of the enzyme’s activity as it was previously shown that most plant and bacterial PMEs have a neutral to alkaline pI, while fungal enzymes are acidic (Sénéchal et al., 2014b). AtPME2 is inhibited *in vitro* by both pH-sensitive and pH-insensitive PMEIs (AtPMEI4 (data not shown) and AtPMEI9 (**Figure 3A**), respectively) (Hocq, Sénéchal, et al., 2017). This suggests that it might be the target of multiple inhibitors at the cell wall, which further questions the role of such diversity of PME-PMEI interactions.

We applied a newly developed LC-MS/MS oligosaccharide-profiling approach, that determines DP and methylation of OGs, for analysing PME processivity (Voxeur et al., 2019; Hocq et al., 2020). We determined that AtPME2 presents a non-processive mode of action at pH 5, close to cell wall pH, while it is processive at pH8, close to its optimum of activity (Hocq et al., 2017). Processive behaviour of plant and bacterial PMEs have been described in details (Willats et al., 2006; Jolie et al., 2010), and the structural determinants of the processivity of the *Erwinia* PME was unraveled using molecular dynamic (MD) simulations (Mercadante et al., 2013; Mercadante et al., 2014), where the rotation of monosaccharide subunits in the binding groove of the enzyme was shown to be a key determinant of the processivity. In contrast to plant and bacterial PMEs, fungal enzymes are often regarded as non-processive. The elucidation of the 3D structure of a non-processive, salt-requiring, PME from *Aspergillus niger* demonstrated key differences between processive and non-processive isoforms, highlighting the importance of the electrostatic potentials of the enzymes in determining their processivity (Kent et al., 2016). This is in accordance with our experimental observations of the differences in processivity observed at different pH for the plant PMEs. Differences in the electrostatic similarity indices (Blomberg et al., 1999; Wade et al., 2001) calculated at two different pH can therefore yield an understanding of the different properties of PMEs at different pH. Interestingly, the comparison of the electrostatic potentials of PMEs either active in basic or acidic conditions suggests that differences between the electrostatic potentials at acidic and alkaline pH are particularly concentrated in proximity of subsites −2 and −3, which have been identified as preferentially docking negatively charged de-methylesterified galacturonic acid monomers; non-processive AnPME show the absence of positive patches in these subsites. Our results clearly demonstrate that the pH-dependency of the mode of action of PMEs, previously suggested in early reports on apple PME (Catoire et al., 1998), might be a key to fine-tune enzymes activity in cell wall microenvironments defined by local pH. Localized changes in apoplastic pH were indeed previously shown to be of major importance for auxin-mediated hypocotyl elongation (Fendrych et al., 2016). As such, pH-dependent changes in PME mode of action might explain a number of unexpected results linking pectin chemistry to cell wall mechanical properties gathered over the last 10 years. Based on *in vitro* studies on pectin-based gels, it was assumed that demethylesterification of pectins by plant PME should lead to large stretches of negatively-charged GalA that can cross-link with calcium ions, stiffening the wall (Willats et al., 2001). However, this scheme appears contradictory with reports showing that overexpressing plant PMEs lead to reduced stiffness of the wall, through decreased values of the Young’s modulus (Peaucelle et al., 2011a; Peaucelle et al., 2015; Wang et al., 2020). Supported by our results on AtPME2, a possible explanation of these apparent contradictory reports might reside in the fact that, within the acidic context of the cell wall, PME mode of action is not what was inferred from *in vitro* studies and may change according to pH microenvironments.

To assess whether the absence of AtPME2 can have consequences on development, we analyzed two T-DNA alleles, *pme2-1* and *pme2-2*, knock-out at the gene and protein levels. We showed that *pme2* mutants had reduced root length as well as reduced lateral roots density, compared to the control. Dark-grown hypocotyls of *pme2* mutants were shorter compared to wild-type, which correlated with an increase in cell wall’s Young’s Modulus in elongating cells of this organ, at the top of the hypocotyl. In addition, these mutants showed decreased PME as well as reduced PG activities. It therefore appears that, in *pme2* mutants, higher methylesterification of pectins would prevent their hydrolysis by PGs, reducing elongation through stiffening the walls. Our results are in accordance with those presented by Peaucelle et al. (Peaucelle et al., 2011; Peaucelle et al., 2015), either in elongating organs such as our model, or in meristems, where softening of the wall is a prerequisite for organ initiation. In both cases, softening of the wall was correlated with higher 2F4 labelling (Braybrook and Peaucelle, 2013; Peaucelle et al., 2015).

## Conclusions

According to our results, in the acidic context of the cell wall, AtPME2 would participate to pectin demethylesterification by randomly acting on the HG chains, leading to the creation of substrates for PG, and consequent destructuration of HG. The somehow contrasting reports linking pectins to cell wall mechanics (Peaucelle et al., 2015; Daher et al., 2018) might be partially explained by our results, suggesting that pH plays a key role in changing PME processivity, thereby affecting pectin mechanical properties. Our biochemical data support a model for which the regulation of PME activity by microdomains of distinct pH might be a key to link pectin chemistry to cell wall mechanics.

## Material and Methods

### Plant material and growth conditions

Two *Arabidopsis thaliana* homozygous T-DNA insertion lines for *At1g53830* (*PME2*) were selected by PCR (see primers in **Supplemental Table II**): *pme2-1* is in Col-0 ecotype (, GK-835A09, in the third exon) and and *pme2-2* in WS ecotype (FLAG445B05, in the first exon). For RT-qPCR analysis of *AtPME2* gene expression, seeds from Columbia-0 (Col-0) background were sowed either on soil or on plates containing ½ MS solid media and grown in light and dark conditions as previously described (Hocq et al., 2020). For kinetic phenotyping of hypocotyls, seeds from the four genotypes were sterilized and sown *in vitro* (Hocq et al., 2020). They were then put in the dark at 21°C for 6 days in a phenobox chamber. This specific growth chamber is designed to receive 27 square Petri dishes (12cm*12cm) and to allow automatic image acquisition of each one using a 36 Mpix D810 camera (Nikon, Champigny sur Marne, France) fixed onto a robotic arm (Optimalog, Saint-Cyr-sur-Loire, France). Pictures were taken every 4 hours, during the first 20 hours, and then every 2 hours. Images from each seedling were analyzed by a specific software (Optimalog) for measurement of hypocotyl length. For each biological replicate, at least 40 hypocotyls were analyzed.

### Analysis of gene expression by RT-qPCR

Total RNAs extraction, cDNA synthesis and RT-qPCR experiments were performed as previously described (Hocq et al., 2020) using specific primers for *AtPME2* (**Supplemental Table II**) Relative expression was normalized according to the most stable reference genes, identified with Genorm in each sample panels (Vandesompele et al., 2002): *CLA* (*At5g46630*, for different organs) and *TIP41* (At4G34270 for dark-grown hypocotyl development. Method used to determine relative expression was previously described.(Sénéchal et al., 2014). Two to three biological replicates were realized, with two technical replicates each.

### Promoter amplification, plant transformation and GUS staining

Amplification of the promoter sequence of At *PME2* (~2 kb upstream of the *AtPME2* transcription start) was performed using the specific primers (**Supplemental Table II**). The purified PCR product was subsequently cloned into pBI101.3 (Ozyme, Saint-Cyr l’Ecole, France), upstream of *GUS* coding gene. Transformation, plant selection, GUS staining and image acquisition was as previously described (Hocq et al., 2020).

### Fusion of *AtPME2* coding sequence with fluorescent tag and confocal imaging

*At1g53830* coding sequence was amplified from Riken pda01692 by PCR using Phusion Hot Start II DNA Polymerase (Thermo Scientific, F549) and specific primers (**Supplemental Table II**). PCR product was fused to GFP CDS in pGBW454 under the control of CaMV-P35S promoter using LR cloning. *Rhizobium radiobacter* (C58C1) was transformed with pGBW405 recombinant vector *via* electroporation and used for transformation of Arabidopsis. After selection of transformants, roots were incubated with propodium iodide (IP, 0.1 mg ml^−1^, Sigma-Aldrich, # P4864, St. Louis, MO, USA) for 20 minutes, transferred in 1 M sorbitol solution to plasmolyze cells before observation under confocal microscope (Zeiss, LMS 780). Excitation wavelengthes are 370-560 nm and 488 nm for IP and GFP respectively, and emission wavelengthes are 631 nm and 493-549 nm, respectively.

### AtPME2 cloning and overexpression in *Pichia pastoris* and purification

For cloning in expression vector, the coding sequence, minus the signal peptide, of *At1g53830* was amplified from Riken pda01692, using the Phusion Hot Start II DNA Polymerase and specific primers (**Supplemental Table II**) The amplified full-length sequence, referred as “FL”, was cloned in frame with polyHis sequence into pPICZαB (ThermoFisher Invitrogen™) as previously described (Hocq *et al*. 2020). Transformation, selection of transformants and cultures were performed as previously described (Hocq et al., 2020). Culture supernatants recovered following centrifugation, were applied onto a CM-FF Hi trap cation-exchange column following the manufacturer’s instructions (GE-Healthcare). Fractions with PME activity were pooled and concentrated. For LC-MS determination of the mode of action, aliquots of AtPME2 were exchanged into 100 mM ammonium acetate pH 5, or 20 mM Tris HCl pH 8 using PD SpinTrap G-25 (GE-Healthcare, 28-9180-04) following manufacturer’s recommendations.

### Mass spectrometry analysis of PMEs

Cell wall-enriched protein fractions from 4 day-old dark-grown hypocotyls (WT and *pme2* mutants) and roots (WT) were extracted from 50 mg frozen fine powder according to method previously described (Sénéchal et al., 2014a). Equal amounts of proteins were resolved on SDS-PAGE for each condition. Tryptic peptides from excised bands were separated and analysed as previously described (Sénéchal et al., 2014a). Following purification of AtPME2 by cation exchange chromatography, bands corresponding to putative mature and PRO part were excised and treated as described above.

### PME-specific antibodies and Western blot analysis

For Western blot analysis of recombinant AtPME2, AtPME3 (Sénéchal et al., 2015), purified native sweet orange and tomato PMEs (Savary, 2001; Savary et al., 2010; Savary et al., 2013), and cell wall-enriched protein extracts from dark-grown hypocotyls and roots, were separated onto a SDS-PAGE and proteins were transferred to Hybond-P PVDF transfer membrane (GE Healthcare, Amersham™ RPN303F) using the manufacturer’s instructions and a Trans-Blot TURBO Transfer System (Bio-Rad, 170-4155) at 0.1A for 30 min. Blotted membranes were blocked with BSA and incubated for 2 h at room temperature with 1:3000 dilution of anti-PME primary antibody. This polyclonal antibody was raised in rabbits against a synthetic peptide (CKTYLGRPWKEYSRT) (Genscript, Piscataway, NJ, USA) that includes the highly conserved amino acid sequence including residue in the catalytic site of PMEs (Markovič and Janeček, 2004). Blotted membrane was probed with 1:5000 dilution of anti-rabbit secondary antibody coupled with peroxidase (ThermoFisher, 31460), followed by detection with the chemiluminescent substrate (ECL™ Prime Western Blotting System, GE Healthcare, RPN2232).

### PME activity assays

Total PME activity was quantified on cell wall-enriched protein extracts using commercial citrus pectins (DM >85% P9561, Sigma-Aldrich) and the alcohol oxidase-coupled colorimetric assay (Klavons and Bennet, 1986; L’Enfant et al., 2015). Substrate specificity of recombinant AtPME2 activity was determined at pH 7.5 and 28°C using commercial citrus pectin (Sigma-Aldrich, DM >85%, P9561; DM 55-70, P9436; DM 20-34%, P9311), sugar beet pectin (DM 42%, degree of acetylation 31% (CPKelco). Results were expressed as nmol MeOH min^−1^ μg^−1^ of protein using a methanol standard curve. The kinetic parameters, V_max_ and K_m_, were determined on citrus pectin (DM 55-70%, Sigma-Aldrich, P9436). The reactions were performed with 3 to 6 replicates using substrate concentrations ranging from 0.125 to 2 mg mL^−1^. The kinetic data were calculated by the Hanes-Wolf plot. Total PG activity from cell wall enriched dark-grown hypocotyl extract was determined as previously described (Hocq et al., 2020). The effects of pH on purified AtPME2 activity; the inhibition assays with PMEI9 were quantified by gel diffusion (Downie et al., 1998) with some modifications (Ren and Kermode, 2000).

### Oligosaccharide oligoprofiling

To determine the mode of action of recombinant AtPME2, first, 0.4% (w:v) citrus pectin DM 55-70 % (Sigma, Cat. No. P9436) were first subjected to a 2 h digestion in ammonium acetate buffer 100mM pH4 at 40 °C by 2.9 U.mL^−1^ of *Aspergillus acuelatus endo*-polygalacturonase M2 (Megazyme, Bray, Ireland) to generate OGs which differ in their degrees of polymerization and methylesterification. After addition of 1 volume absolute ethanol and centrifugation (5 min, 5000g), the upper phase containing OGs was divided in two tubes, evaporated, and re-suspended either in ammonium acetate 100mM pH 5, or in Tris HCl 20mM pH 8. After heat-inactivation of PG, OGs were treated for 16h at 40°C by 80 nmol/min of purified AtPME2 either at pH 5 (Buffer) or 20 nmol/min at pH 8 (Buffer) in order to compensate for the difference in activity measured at both pH. Chromatographic separation of OGs by size exclusion chromatography (SEC), MS-detection and data acquisition and processing were performed as previously described (Hocq et al., 2020).

### Calculation and comparison of protein electrostatics

3D models of putative mature parts of Arabidopsis AtPME2 and Orange PME (CsPME4) were created using the I-TASSER prediction software (Zheng et al., 2019), with *D. carota* PME (PDB:1GQ8), (Johansson et al., 2002) as starting model. Protein electrostatic potentials were obtained by solving the linearized version of the Poisson-Boltzmann (PB) equation using APBS version 3.0 (Baker et al., 2001). Atomic radii and partial charges were assigned according to the AMBER99 force field parameters (Wang et al., 2000), using PDB2PQR version 2.1.1 (Dolinsky et al., 2004). The structures of AtPME2, AnPME (PDB: 5C1C) and CsPME4 were protonated considering the protonation states empirically estimated using PROPKA version 3.3 (Søndergaard et al., 2011) at either pH 5.0 or pH 8.0. The solution of the PB equation was discretized on a 19.3 nm^3^ grid with spacing of 0.6 Å, centered on the C_α_ atom of one of the PMEs catalytic aspartic acid residues, conserved across PMEs. Solvent dielectric was set at a value of 78.5 to account for an aqueous environment, whereas solute dielectric and temperature were set to 4.0 and 298.15 K respectively. Potentials calculated at pH 8.0 were then subtracted, grid point by grid point, from the potentials calculated at pH 5.0 to obtain an electrostatic potential difference. A numerical comparison of the electrostatic potentials was achieved by calculating electrostatic similarity indices as the cross-product between two electrostatic potentials:

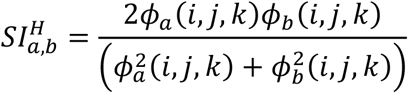

Where 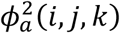 and 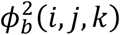 are the electrostatic potentials calculated at the grid points *i,j,k* for proteins *a* and *b* (Blomberg et al., 1999; Wade et al., 2001).

### Atomic Force Microscopy (AFM) measurements

The protocol was adapted from (Milani et al., 2011) and applied to wild type and *pme2-1* hypocotyls at 3 days after induction of germination. Hypocotyls were immobilized and covered with water. Measurements were performed as close as possible (typically ~ 1 mm) to the hook (**Supplemental Figure 6A**). We used cantilevers with pyramidal tip and spring constant 5-6N/m. We analysed three regions of size 60μmx60μm per hypocotyl and obtained force-depth curves on ~60 points along the top of visible cells (**Supplemental Figure 6B**), with maximal depth 100-400 nm. Apparent Young’s modulus was obtained by fitting the 0-100 μm depth range of the force-depth curve with the Sneddon model for a cone of half-angle 18°, assuming a Poisson ratio of 0.3 for the cell wall (**Supplemental Figure 6C)**. We pooled together all values of modulus for a given hypocotyl.

### Statistical analysis

Data represent the mean ± SE and were treated with R software (R development Core Team, 2008). Normality of data and equality of variances were assessed using Shapiro-Wilk and F-tests, respectively. Non-parametric Wilcoxon test was carried out for pairwise comparisons. Significant differences between two groups were determined as highly significant for p < 0.001 (***), significant for p < 0.1 (**), and moderately significant for p < 0.05 (*), while ns indicates non-significant differences.

## Accession numbers

Nucleotide sequence data from this article can be found in The Arabidopsis Information Ressource database under the following accession numbers : *AtPME2, AT1G53830 ; AtAtPMEI9, AT1G62770 ; CLA, AT5G46630 ; APT1, AT1G27450 ; TIP41, AT4G34270.* Protein data from this article can be found in UniprotKB database under the following accession numbers: AtPME2, Q42534; AtPME3, O49006 ; AtPMEI9, Q9SI72.

## Notes

**Funding information** This work was supported by grants from the Agence Nationale de la Recherche (ANR-12-BSV5-0001 GALAPAGOS and ANR PECTOSIGN). LH was recipient of a studentship from the “Trans Channel Wallnet” project, which was selected by the INTERREG IVA program France (Channel) – England European cross-border cooperation program. The financial support from the Institut Universitaire de France (IUF) to JP is gratefully acknowledged.

### Competing Interest Statement

The authors have declared no competing interest.

