## supplemental figures for "Arabidopsis AtPME2 has a pH-dependent processivity and control cell wall mechanical properties"

```
  1 MAPIKEFISK FSDFKNNKKL ILSSAAIALL LLASIVGIAA TTTNQNKNQK
                                             FL
 51 ITTLSSTSHA ILKSVCSSTL YPELCFSAVA ATGGKELTSQ KEVIEASLNL
101 TTKAVKHNYF AVKKLIAKRK GLTPREVTAL HDCLETIDET LDELHVAVED
151 LHQYPKQKSL RKHADDLKTL ISSAITNQGT CLDGFSYDDA DRKVRKALLK
201 GQVHVEHMCS NALAMIKNMT ETDIANFELR DKSSTFTNNN NRKLKEVTGD
251 LDSDGWPKWL SVGDRRLLQG STIKADATVA DDGSGDFTTV AAAVAAAPEK
                              ****** ********** **********
301 SNKRFVIHIK AGVYRENVEV TKKKTNIMFL GDGRGKTIIT GSRNVVDGST
                    ***** ***
351 TFHSATVAAV GERFLARDIT FQNTAGPSKH QAVALRVGSD FSAFYQCDMF
401 AYQDTLYVHS NRQFFVKCHI TGTVDFIFGN AAAVLQDCDI NARRPNSGQK
451 NMVTAQGRSD PNQNTGIVIQ NCRIGGTSDL LAVKGTFPTY LGRPWKEYSR
            ** ********** ***            ****** ******
                             ******* ****
501 TVIMQSDISD VIRPEGWHEW SGSFALDTLT YREYLNRGGG AGTANRVKWK
551 GYKVITSDTE AQPFTAGQFI GGGGWLASTG FPFSLSL*
```

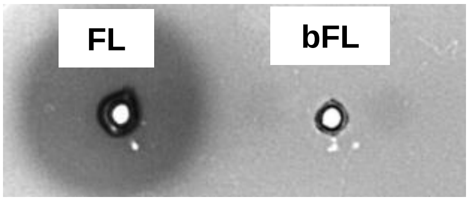

**Supplemental Figure 1:** AtPME2 amino-acid sequence highlighting the sequence used for expression in *Pichia pastoris* (FL, PRO part in green, mature part in cyan and putative processing motifs indicated in red). Peptides mapping the mature part of AtPME2 in cell wall-enriched extracts of Col-0 dark-grown hypocotyls using nano LC-MS/MS are underlined with asterisks.

```
  1 MAPIKEFISK FSDFKNNKKL ILSSAAIALL LLASIVGIAA TTTNQNKNQK
 51 ITTLSSTSHA ILKSVCSSTL YPELCFSAVA ATGGKELTSQ KEVIEASLNL
101 TTKAVKHNYF AVKKLIAKRK GLTPREVTAL HDCLETIDET LDELHVAVED
151 LHQYPKQKSL RKHADDLKTL ISSAITNQGT CLDGFSYDDA DRKVRKALLK
201 GQVHVEHMCS NALAMIKNMT ETDIANFELR DKSSTFTNNN NRKLKEVTGD
251 LDSDGWPKWL SVGDRRLL
                      QG STIKADATVA DDGSGDFTTV AAAVAAAPEK
301 SNKRFVIHIK AGVYRENVEV TKKKTNIMFL GDGRGKTIIT GSRNVVDGST
351 TFHSATVAAV GERFLARDIT FQNTAGPSKH QAVALRVGSD FSAFYQCDMF
401 AYQDTLYVHS NRQFFVKCHI TGTVDFIFGN AAAVLQDCDI NARRPNSGQK
451 NMVTAQGRSD PNQNTGIVIQ NCRIGGTSDL LAVKGTFPTY LGRPWKEYSR
501 TVIMQSDISD VIRPEGWHEW SGSFALDTLT YREYLNRGGG AGTANRVKWK
551 GYKVITSDTE AQPFTAGQFI GGGGWLASTG FPFSLSL
```

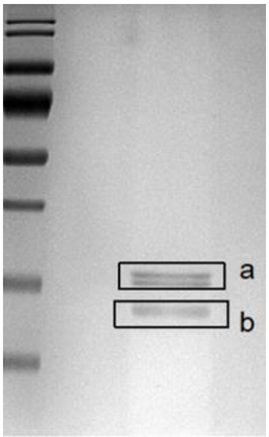

**<u>Catalytic region tryptic peptides – 12 observed (2 truncated) (“Band a”)</u>**

| **Position** | **Peptide sequences** |
|---|---|
| 75-300 | ADATVADDGSGDFTTVAAAVAAAPEK* |
| 316-322 | ENVEVTK* |
| 325-334 | TNIMFLGDGR |
| 344-363 | NVVDGSTTFHSATVAAVGER |
| 368-379 | DITFQNTAGPSK |
| 387-412 | VGSDFSAFYQCDMFAYQDTL YVHSNR |
| 451-458 | NMVTAQGR |
| 459-473 | SDPNQNTGIVIQNCR* |
| 474-484 | IGGTSDLLAVK* |
| 485-496 | GTFPTYLGRPWK* |
| 501-532 | TVIMQSDISDVIRPEGWH (+EWSGSFALDTLTYR) – fragment observed |
| 554-587 | VITSDTEAQPFTAGQFIGGG (+GWLASTGFPFSLSL) – fragment observed |

**<u>PRO-region tryptic peptides – 7 observed (“Band b”)</u>**

| **Position** | **Peptide sequences** |
|---|---|
| 51-63 | ITTLSSTSHAILK |
| 64-85 | SVCSSTLYPELCFSAVAATGGK |
| 107-113 | HNYFAVK |
| 169-192 | TLISSAITNQGTCLDGFSYDDADR |
| 201-217 | GQVHVEHMCSNALAMIK |
| 218-230 | NMTETDIANFELR |
| 233-242 | SSTFTNNNNR |

*Peptides observed in cell wall extract and indicated by sequence alignment in Supp Fig 1

**Supplemental Figure 2:** Identification of peptides mapping the PRO and mature part of AtPME2 following cation exchange purification. Band **a** corresponds to the two proteins at ~35 kDa while band **b** corresponds to the band below ~35kDa (see gel).

**A**

```
At3g62170           IPDKELEADK----LTIKSYLGRPWKKFATTVIIGTEIGDLIKPEGWTEWQG-EQN--HK
At2g47030           VPDRKLTPER----LTVATYLGRPWKKFSTTVIMSTEMGDLIRPEGWKIWDG-ESF--HK
At2g47040           VPDKKLAAER----LIVESYLGRPWKKFSTTVIINSEIGDVIRPEGWKIWDG-ESF--HK
At2g26450           AANEDLKPVK----EEYKSYLGRPWKNYSRTIIMESKIENVIDPVGWLRWQETDFA--ID
At4g33230           APNEDLKPVK----AQFKSYLGRPWKPHSRTVVMESTIEDVIDPVGWLRWQETDFA--ID
At3g10710           SPLGDLTD--------VMTFLGRPWKNFSTTVIMDSYLHGFIDRKGWLPWTG-DSA--PD
At5g04960           KPLDNLTD--------IQTFLGRPWKDFSTTVIMKSFMDKFINPKGWLPWTG-DTA--PD
At3g10720           KAAPDLAAEP----KSAMTFLGRPWKPYSRTVFMQSYISDIVQPVGWLEWNG-TIG--LD
At5g04970           GAAPDLAADP----KSTMTFLGRPWKPYSRTVYIQSYISDVVQPVGWLEWNG-TTG--LD
At4g03930           TASSDLDTAT------VKTYLGRPWRIFSTVAVLQSFIGDLVDPAGWTPWEG-ETG--LS
At1g11590           TASSDLDTTT------VKTYLGRPWRIFSTVAVMQSFIGDLVDPAGWTPWEG-ETG--LS
At3g27980           TTSSDLDTAT------VKTYLGRPWRRFSTVAVLQSFIGDLVDPAGWTPWKG-ETG--LS
At5g20860           RTDSDLSPVK----HKYSSYLGRPWRKYSRAIVMESYIDDAIAEGGWAGWLD-SGDEVLK
At5g51490           LPAPDLKPVV----GTVKTYMGRPWMKFSRTVVLQTYLDNVVSPVGWSPWIE-GSVFGLD
At5g51500           IPAPDLKPVV----RSVKTYMGRPWMMYSRTVVLKTYIDSVVSPVGWSPWTK-GSTYGLD
At4g15980           TGDASYLPVK----AKNRAFLGRPWKEFSRTIIMNTEIDDVIDPEGWLKWNE-TFA--LN
At3g47400           IAASDLKPVI----RAYKTYLGRPWQAYSRVTIMKTYIDNSISPLGWSPWLR-GSNFALN
At3g14300           SPNGNVT---------ATTYLGRPWKLFSKTVIMQSVIGSFVNPAGWIAWNS-TYDPPPR
At1g23200           ATA-------------SETYLGRPWRSHSRTVFMKCNLGALVSPAGWLPWSG-SFA--LS
At5g64640           NGTEEYMKEFQANPEGHKNFLGRPWKEFSRTVFVNCNLESLISPDGWMPWNG-DFA--LK
At5g53370           LATPDLEASK----GSYPTYLGRPWKLYSRVVYMMSDMGDHIDPRGWLEWNG-PFA--LD
At5g49180           TGEPAYIPVK----SINKAYLGRPWKEFSRTIIMGTTIDDVIDPAGWLPWNG-DFA--LN
At5g09760           NGTEEYMKLFKANPKVHKNFLGRPWKDYSRTVFIGCNLEALITPDGWLPWSG-DFA--LK
At4g02320           LAAPDLIPVQ----ANFKAYLGRPWQLYSRTVIMKSFIDDLVDPAGWLKWKD-DFA--LE
At4g02300           LAAPDLIPVK----ENFKAYLGRPWRKYSRTVIIKSFIDDLIHPAGWLEGKK-DFA--LE
At4g00190           KGAPGVQ--L----GGVKTYLGRPWRSYARTVVIGTYLDTLIEPNGWIDWDN-VTA--LS
At3g49220           LAASDLQATN----GSTQTYLGRPWKLFSRTVYMMSYIGGHVHTRGWLEWNT-TFA--LD
At3g06830           TGDPAYIPMK----SVNKAYLGRPWKEFSRTIIMKTTIDDVIDPAGWLPWSG-DFA--LK
At2g26440           LASEDLFNSS----NKVKSYLGRPWREFSRTVVMESYIDEFIDGSGWSKWNG-GEA--LD
At1g53840           SANGNVI---------APTYLGRPWKEFSTTVIMETVIGAVVRPSGWMSWVS-GVDP-PA
At4g33220           SADADLVPYL----NTTRTYLGRPWKLYSRTVFIRNNMSDVVRPEGWLEWNA-DFA--LD
At3g60730           RAAPEFEAVK----GRFKSYLGRPWKKYSRTVFLKTDIDELIDPRGWREWSG-SYA--LS
At3g43270           AADTDLLLNL----NTTATYLGRPWKLYSRTVFMQNYMSDAINPVGWLEWNG-NFA--LD
At3g05620           LAT-------------QPTYLGRPWKLYSRTVYMNTYMSQLVQPRGWLEWFG-NFA--LD
At3g05610           AGEPDYLAVK----ETSKAYLGRPWKEYSRTIIMNTFIPDFVQPQGWQPWLG-DFG--LK
At2g45220           TAASDLRPVL----GSTKTYLGRPWRQYSRTVFMKTSLDSLIDPRGWLEWDG-NFA--LK
At1g11580           TASSDLAPVK----GSVKTFLGRPWKLYSRTVIMQSFIDNHIDPAGWFPWDG-EFA--LS
➡ At3g14310         GATSDLQSVK----GSFPTYLGRPWKEYSQTVIMQSAISDVIRPEGWSEWTG-TFA--LN
At5g27870           VGEPDYLAVK----EQSKTYLGRPWKEYSRTIIMNTFIPDFVPPEGWQPWLG-EFG--LN
At3g59010           TGS-------------TKTYLGRPWKQYSRTVVMQSFIDGSIHPSGWSPWSS-NFA--LK
At2g43050           TAE-------------SMTYLGRPWKEYSRTVVMQSFIGGSIHPSGWSPWSG-GFG--LK
➡ At1g53830         GGTSDLLAVK----GTFPTYLGRPWKEYSRTVIMQSDISDVIRPEGWHEWSG-SFA--LD
At4g02330           KPADDLVSSN----YTVKTYLGRPWKEYSRTVFMQSYIDEVVEPVGWREWNG-DFA--LS
At2g47550           RPADDLATSN----YTVKTYLGRPWKEYSRTVVMQTYIDGFLEPSGWNAWSG-DFA--LS
At1g02810           KPADDLVSSN----YTVKTYLGRPWKEYSRTVYMQSYIDGFVEPVGWREWNG-DFA--LS
Antibody            -----------------KTYLGRPWKEYSRTV----------------------------
                                       ::****  .: .
```

**B**

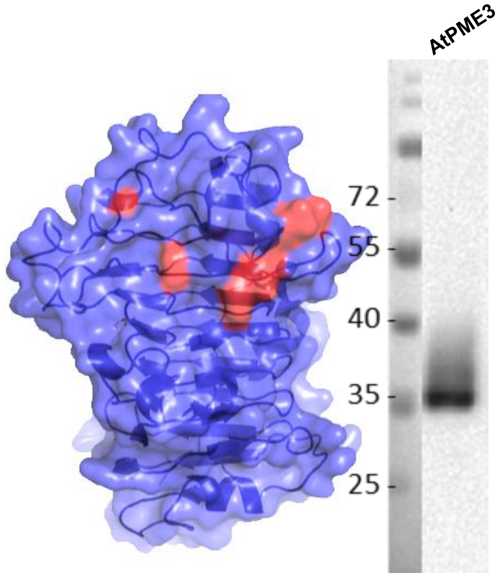

**Supplemental Figure 3: (A)** Sequence alignment of 45 Arabidopsis PME isoforms showing the conservation of the epitope used for production of the generic anti-PME antibody. **(B)** Immunodetection of AtPME3 using the generic anti-PME antibody.

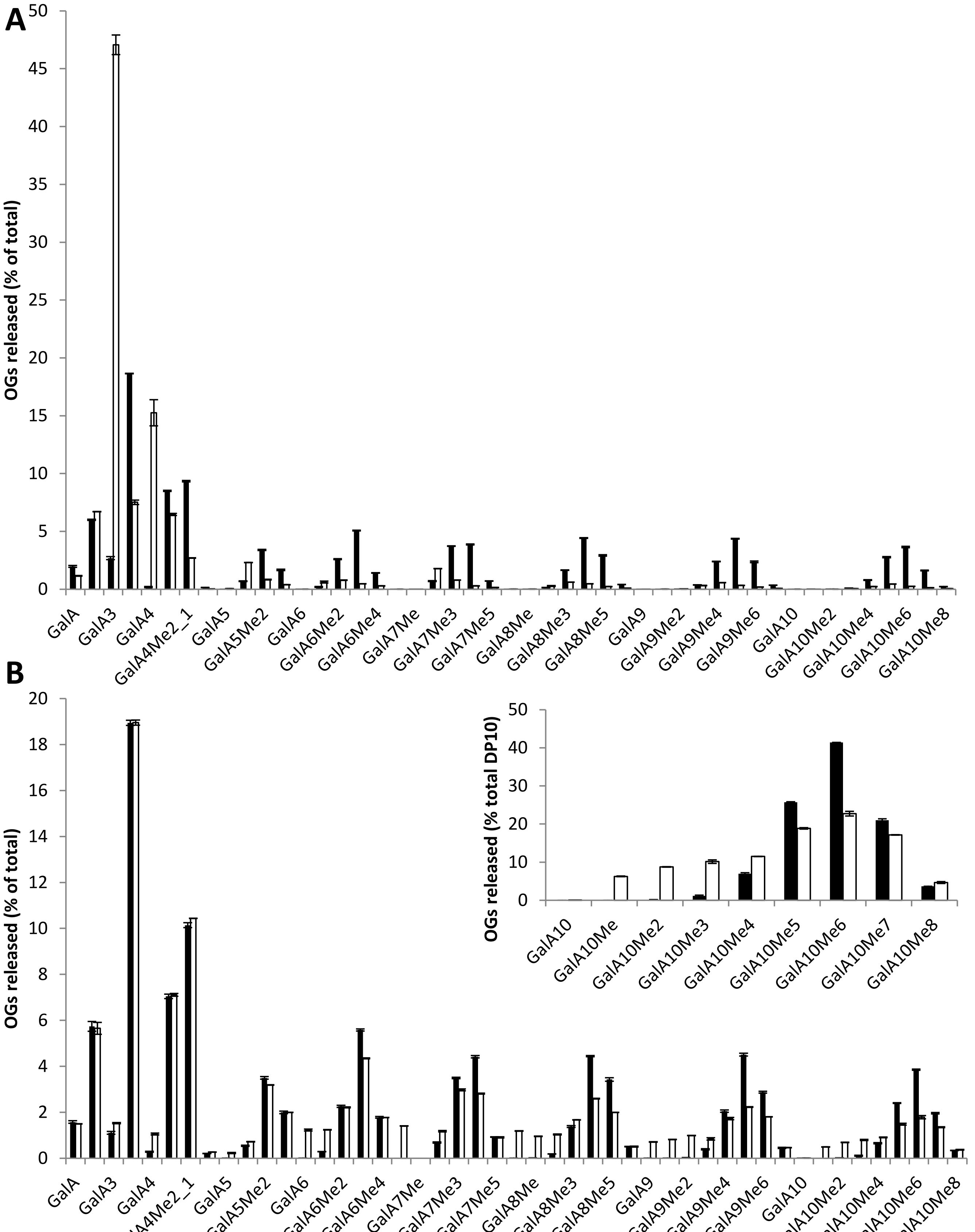

**Supplemental Figure 4:** Determination of the processivity of CsPME at **(A)** pH 8 and (**B**) pH 5 using LC-MS/MS. Non-digested samples (black bars); CsPME digestion (white bars). Data represent the mean ± SE of three replicates.

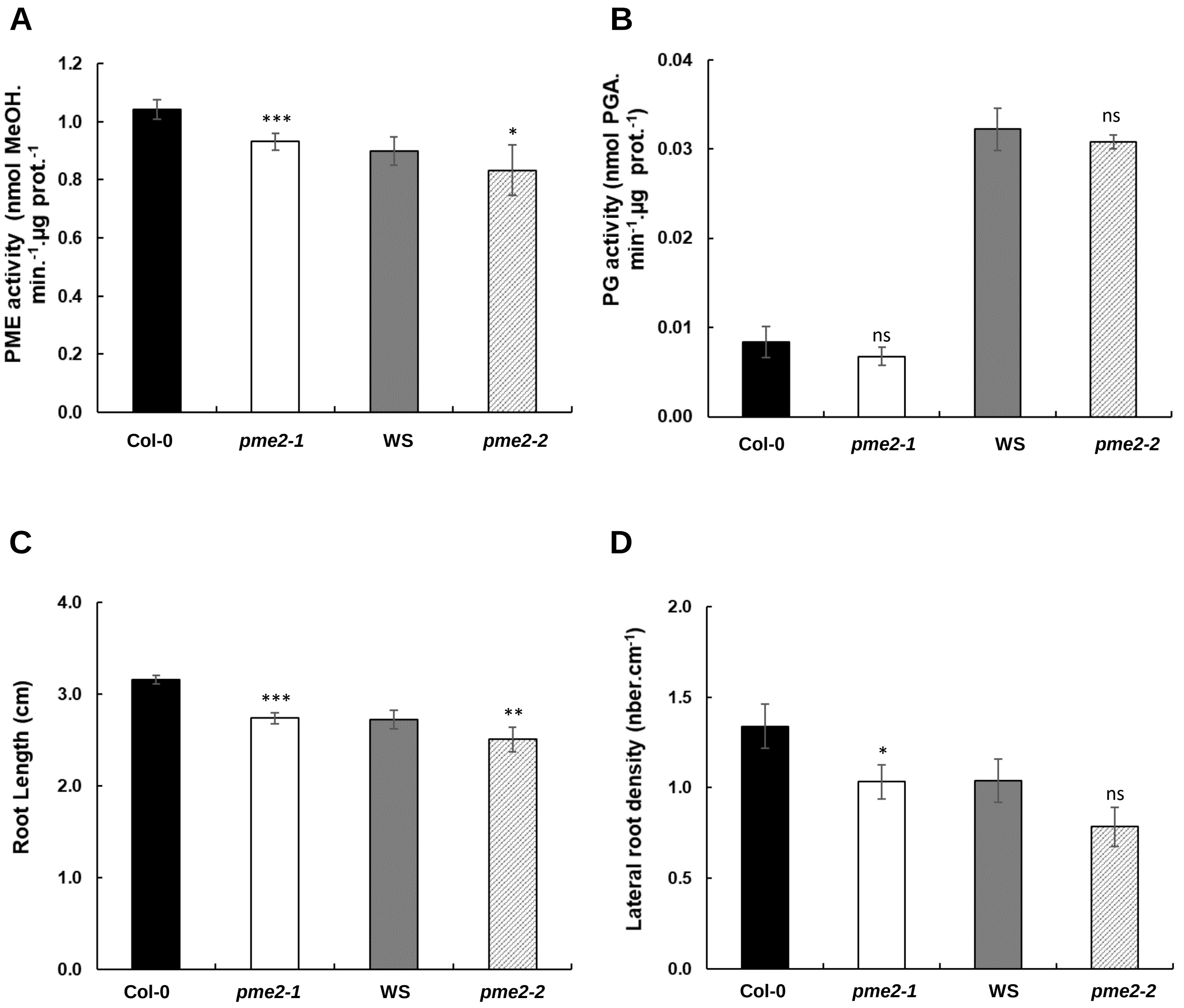

**Supplemental Figure 5:** *pme2* mutation affect root development. **(A**) Total PME activity of cell wall-enriched protein extracts from 7-day-old roots of wild type Col-0/WS and *pme2-1/pme2-2* mutants. Data represent the means of PME activity in nmol of methanol.min$^{-1}$/µg of protein$^{-1}$± SE of three independent protein extractions and three technical replicates (n=9). (**B**) Total PG activity of cell wall-enriched protein extracts from 4 day-old dark-grown hypocotyls of wild type Col-0/WS and *pme2-1/pme2-2* mutants. Data represent the means of PG activity in nmol of PGA.min$^{-1}$/µg of protein$^{-1}$ ± SE of three independent protein extractions and three technical replicates (n=9). **(C)** Root length measured on 7-day-old roots of wild-type and *pme2-1* and *pme2-2* mutant lines (n>90). **(D)** Lateral root density (number of emerged lateral root.cm-1) of wild-type and *pme2-1* and *pme2-2* mutant lines (n>90). Statistical analyses were realized using Mann-Whitney test: * *P*<0.05.

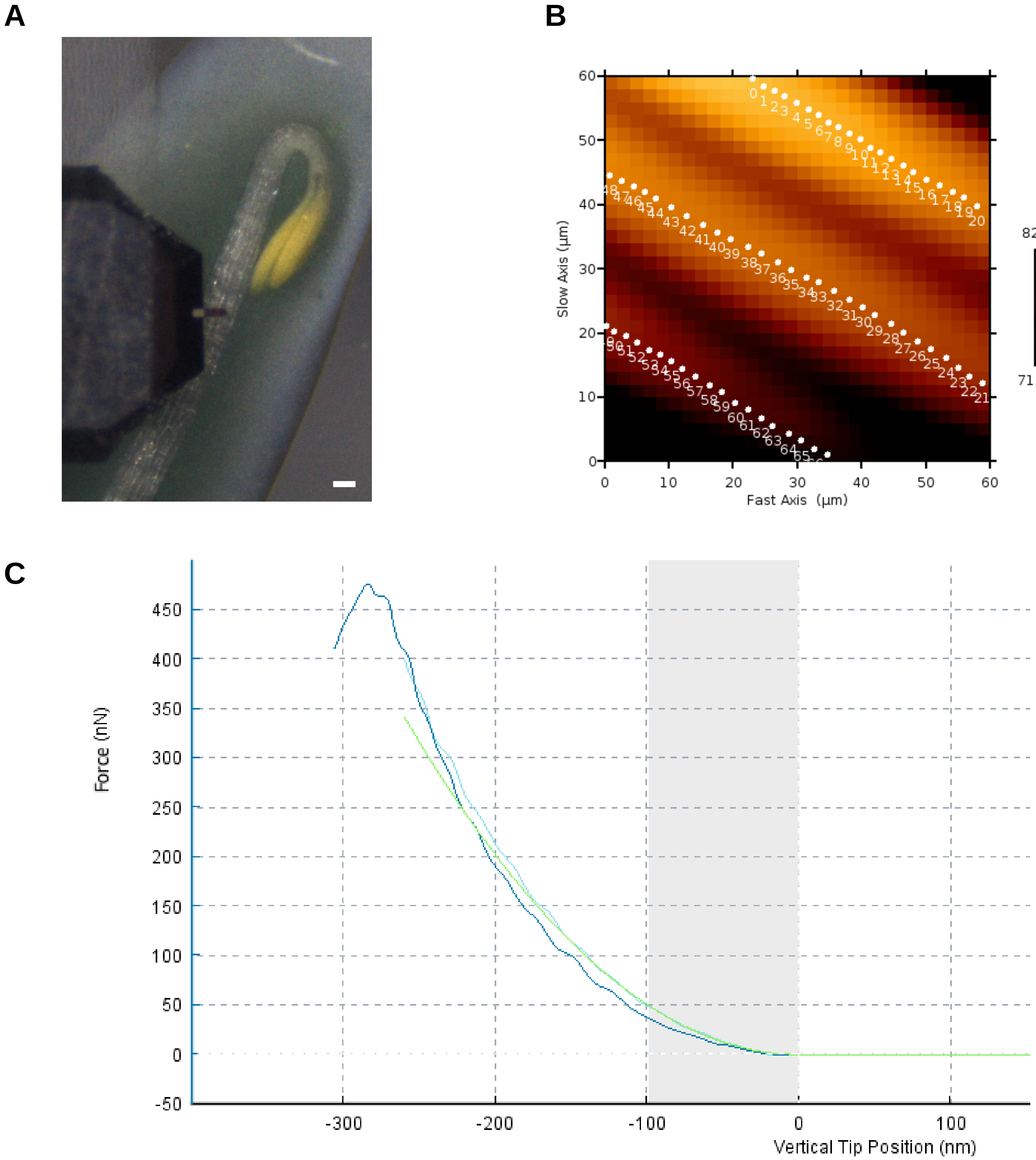

**Supplemental Figure 6:** AFM analysis of 3-day-old hypocotyls. **(A)** Top view of a wild-type hypocotyl under the atomic force microscope; the cantilever (rectangular with a triangular end) is located over the hypocotyl at about 1mm from the hook. Scale bar 100 µm. **(B)** Topographic image of a 60 µm x 60 µm region of the hypocotyl (height scale on right), with three cells partially visible. The white dots show points where force-depth curves were obtained in order to characterise cell wall stiffness. **(C)** Typical force-depth curve with approach in light blue, retract in dark blue. The green curve is a fit of the approach to the Sneddon model, which yields the apparent Young’s modulus.

**A**

| | | Col | | *pme2.1* | | WS | | *pme2.2* | |
|---|---|---|---|---|---|---|---|---|---|
| | | % coverage | Nb peptides | % coverage | Nb peptides | % coverage | Nb peptides | % coverage | Nb peptides |
| AT1G01900.1 | Subtilase family protein | 18.6 | 11 | 14.08 | 8 | 10.85 | 5 | 13.05 | 7 |
| AT1G20160.2 | Subtilisin-like serine endopeptidase protein | 12.88 | 8 | 15.34 | 9 | 14.25 | 9 | 14.25 | 9 |
| **AT1G53830.1** | **Pectin methylesterase 2** | **19.08** | **5** | | | **18.57** | **5** | | |
| AT2G05920.1 | Subtilase family protein | 26.92 | 16 | 31.03 | 17 | 32.36 | 17 | 31.03 | 16 |
| AT2G43050.1 | Plant invertase/pectin methylesterase inhibitor | 25.68 | 10 | 27.22 | 11 | 22.78 | 9 | 18.73 | 7 |
| AT2G46930.1 | Pectinacetylesterase family protein | 17.55 | 5 | 29.09 | 8 | 31.49 | 9 | 31.01 | 8 |
| AT3G05910.1 | Pectinacetylesterase family protein | 27.95 | 7 | 5.54 | 2 | 35.42 | 9 | 26.75 | 7 |
| AT3G06770.2 | Pectin lyase-like superfamily protein | 15.02 | 5 | 8.22 | 2 | 10.34 | 3 | 12.47 | 4 |
| AT3G07010.1 | Pectin lyase-like superfamily protein | 10.82 | 4 | 10.1 | 3 | | | | |
| AT3G09410.3 | Pectinacetylesterase family protein | | | | | 6.06 | 2 | 8.67 | 3 |
| AT3G14067.1 | Subtilase family protein | 24.45 | 12 | 24.45 | 12 | 26 | 12 | 22.91 | 11 |
| AT3G14310.1 | Pectin methylesterase 3 | 27.87 | 11 | 26.52 | 13 | 28.04 | 11 | 28.04 | 14 |
| AT3G16850.1 | Pectin lyase-like superfamily protein | 28.57 | 7 | 32.75 | 9 | 28.57 | 8 | 30.33 | 8 |
| AT3G43270.1 | Plant invertase/pectin methylesterase inhibitor | 15.37 | 5 | 13.47 | 4 | 13.47 | 4 | 11.2 | 3 |
| AT3G49220.1 | Plant invertase/pectin methylesterase inhibitor | 9.2 | 5 | 10.37 | 6 | 12.04 | 7 | 10.87 | 7 |
| AT3G55140.2 | Pectin lyase-like superfamily protein | 7.82 | 2 | 11.4 | 3 | 7.82 | 2 | 10.75 | 3 |
| AT3G57790.1 | Pectin lyase-like superfamily protein | 4.69 | 2 | 10 | 5 | 12.65 | 6 | 10 | 5 |
| AT3G59010.1 | Pectin methylesterase 61 | | | | | 15.88 | 4 | 13.04 | 4 |
| AT3G61490.1 | Pectin lyase-like superfamily protein | | | | | | | 11.97 | 3 |
| AT4G19410.1 | Pectinacetylesterase family protein | 51.41 | 12 | 53.2 | 9 | 58.31 | 13 | 59.85 | 13 |
| AT4G23820.1 | Pectin lyase-like superfamily protein | 32.66 | 10 | 10.14 | 3 | 18.24 | 5 | 9.23 | 2 |
| AT4G25260.1 | Plant invertase/pectin methylesterase inhibitor | 26.87 | 4 | 40.3 | 6 | 32.34 | 6 | 23.38 | 4 |
| AT4G33220.1 | Pectin methylesterase 44 | 19.43 | 6 | 19.24 | 7 | 19.05 | 5 | 16.76 | 5 |
| AT5G20740.1 | Plant invertase/pectin methylesterase inhibitor | | | | | 11.71 | 2 | 11.71 | 2 |
| AT5G45280.2 | Pectinacetylesterase family protein | 71.1 | 16 | 66.24 | 13 | 44.5 | 10 | 44.5 | 10 |
| AT5G46960.1 | Plant invertase/pectin methylesterase inhibitor | | | | | 14.94 | 2 | 14.94 | 2 |
| AT5G51750.1 | Subtilase 1.3 | 10.13 | 5 | | | 10.26 | 5 | 8.21 | 4 |
| AT5G59090.2 | Subtilase 4.12 | 23.94 | 14 | 25.31 | 15 | 12.45 | 7 | 14.09 | 8 |
| AT5G62350.1 | Plant invertase/pectin methylesterase inhibitor | | | | | 11.39 | 2 | | |
| AT5G67360.1 | Subtilase family protein | 24.7 | 15 | 23.65 | 14 | 41.61 | 20 | 23.65 | 14 |

**B**

| | | Col | | WS | |
|---|---|---|---|---|---|
| | | % coverage | Nb peptides | % coverage | Nb peptides |
| AT1G30600.1 | Subtilase family protein | | | 8.65 | 4 |
| AT1G32940.1 | Subtilase family protein | 20.8 | 14 | | |
| **AT1G53830.1** | **Pectin methylesterase 2** | 18.91 | 5 | 16.87 | 4 |
| AT2G04160.1 | Subtilisin-like serine endopeptidase | 15.16 | 9 | 18.39 | 11 |
| AT2G05920.1 | Subtilase family protein | 62.2 | 29 | 62.2 | 30 |
| AT2G45220.1 | Plant invertase/pectin methylesterase inhibitor | 30.53 | 13 | 31.12 | 13 |
| AT3G14067.1 | Subtilase family protein | 38.87 | 17 | 38.35 | 17 |
| AT3G14310.1 | Pectin methylesterase 3 | 23.99 | 8 | 19.93 | 7 |
| AT3G16850.1 | Pectin lyase-like superfamily protein | 18.46 | 5 | 17.58 | 5 |
| AT3G43270.1 | Plant invertase/pectin methylesterase inhibitor | | | 11.2 | 3 |
| AT3G57790.1 | Pectin lyase-like superfamily protein | | | 14.69 | 7 |
| AT4G19410.1 | Pectinacetylesterase family protein | 63.94 | 14 | 47.39 | 14 |
| AT4G20430.2 | Subtilase family protein | 4.21 | 2 | 4.21 | 2 |
| AT4G21650.1 | Subtilase family protein | | | 11.1 | 6 |
| AT4G23500.1 | Pectin lyase-like superfamily protein | 4.65 | 2 | 12.53 | 4 |
| AT4G25260.1 | Plant invertase/pectin methylesterase inhibitor | | | 11.94 | 2 |
| AT4G33220.1 | pectin methylesterase 44 | | | 13.9 | 3 |
| AT4G34980.1 | Subtilase family protein | 18.85 | 11 | 23.04 | 13 |
| AT5G09760.1 | Plant invertase/pectin methylesterase inhibitor | 13.43 | 5 | 11.8 | 5 |
| AT5G44530.1 | Subtilase family protein | 13.81 | 7 | 9.05 | 4 |
| AT5G45280.2 | Pectinacetylesterase family protein | 47.83 | 10 | 43.22 | 10 |
| AT5G59090.2 | Subtilase 4.12 | 30.23 | 16 | 35.15 | 21 |
| AT5G62350.1 | Plant invertase/pectin methylesterase inhibitor | | | 36.63 | 4 |
| AT5G67360.1 | Subtilase family protein | 32.76 | 17 | 42.8 | 21 |

**Supplemental Table I:** Identification of homogalacturonan remodeling enzymes and regulators in cell wall-enriched protein extracts of **(A)** 4 day-old dark-grown hypocotyls of wild type Col-0/WS and *pme2* mutants and **(B)** Col-0/WS roots.

| | |
|---|---|
| **Genotyping** | |
| *pme2-1* F | 5’-GACGGAAGCGGTGACTTTAC-3’ |
| *pme2-1* R | 5’-AGTGTCCAACGCAAAACTCC-3’ |
| *pme2-2* F | 5’-CTCAACACTATACCCGGA-3’ |
| *pme2-2* R | 5’-ACTTTCCTATCGGCGTCG-3’ |
| GK F | 5’-ATATTGACCATCATACTCATTGC-3’ |
| FLAG F | 5’-CTACAAATTGCCTTTTCTTATCGAC-3’ |
| **RT-qPCR** | |
| AtPME2 F | 5’-TACGACGACGCCGATAGGAAAG-3’ |
| AtPME2 R | 5’-ATGTGCTCCACGTGTACCTGAC-3’ |
| APT1 F | 5’-GAGACATTTTGCGTGGGATT-3’ |
| APT1 R | 5’-CGGGGATTTTAAGTGGAACA-3’ |
| CLA F | 5’-GTTTGGGAGAAGAGCGGTTA-3’ |
| CLA R | 5'-CTGATGTCACTGAACCTGAACTG-3' |
| TIP41 F | 5’-GCTCATCGGTACGCTCTTTT-3’ |
| TIP41 R | 5’-TCCATCAGTCAGAGGCTTCC-3’ |
| **Promoter cloning** | |
| primer F | 5’-**AAGCT**TGTACAATGATGGTTCTATTGT-3’ |
| primer R | 5’-**TCTAGA**GGTGGAATAGGGTTTATATTG-3’ |
| **CDS cloning for confocal imaging** | |
| primer F | 5’-**CACC**ATGGCACCAATCAAAG-3’ |
| primer R | 5’-AAGACTTAACGAGAAAGGAA-3’ |
| **CDS cloning for Pichia expression** | |
| primer F | 5'-**CTGCAG**GAGCCACAACAACAAAC-3' |
| primer R | 5'-**GCGGCCG**CAAGACTTAACGAGAAAGGA-3' |

**Supplemental Table II:** Primers used in the study.
